# Fluorescently guided workflow with rationally engineered 5′ ligation adapters for high-sensitivity and low-bias small RNA sequencing

**DOI:** 10.64898/2026.06.23.733996

**Authors:** Sarah Anne Barnes, Daniel Lovíšek, Natália Dzurčaninová, Martin Čarnecký, Stanislava Bírová, Ivana Čirková, Ján Matyašovský, Adrián Szobi, Pavol Čekan

## Abstract

MicroRNAs (miRNAs) act as key regulators of gene expression across diverse cellular processes, and their precise quantification can provide unique insight into disease pathogenesis. High-throughput sequencing allows for comprehensive small RNA profiling; however, standard commercial library preparation workflows are challenged by issues of low sensitivity and representational bias, limiting reliable profiling, especially in scenarios where samples are scarce. Several structural studies have shown that this bias primarily arises due to sequence and secondary structure variations between miRNAs and adapters during enzyme-catalyzed biochemical reactions. In this work, we propose a new approach to ligation adapter engineering using a bioinformatic analysis of the human miRNome to rationally design structure-forcing 5’ adapters, that physically override localized, unpredictable structural variations during the intermediate ligation state. We show that this approach combined with a practical fluorescence-guided workflow, utilizing a fluorescently-labeled 3’ adapter and novel Fluorescent Ligation Rulers (FLRs) to guide precise band excision, can minimize representational bias and increase the sensitivity of small RNA sequencing from low-input biological matrices. In comprehensive benchmarks using a synthetic panel, this method significantly reduced bias and outperformed alternative commercial protocols. Finally, we demonstrate that this workflow enhances biomarker detection and library quality in challenging clinical matrices, especially in cerebrospinal fluid. Overall, this protocol enables highly accurate miRNome characterization and is well-suited for biomarker discovery in challenging sample types.

## Introduction

MicroRNAs (miRNAs), a class of small non-coding RNAs, act as fundamental regulators of gene expression across diverse cellular processes ^1–3^. Typically spanning 18 to 25 nucleotides in length, these molecules modulate post-transcriptional silencing by binding to target messenger RNAs, thereby controlling translation and transcript stability ^4^. The precise quantification of the small RNA transcriptome can provide unique insight into cellular state, developmental pathways, and disease pathogenesis ^5^. Consequently, small RNA profiling serves as a vital tool for basic biological discovery, the identification of clinical biomarkers, and the characterization of therapeutic targets.

High-throughput sequencing of miRNAs, the most widely used approach to study these potential biomarkers in complex biological samples, necessitates the precise conversion of the native small RNA pool into a sequenceable complementary DNA (cDNA) library. The standard ligation-based library preparation workflow involves a strictly defined sequence of enzyme-catalyzed biochemical reactions. Initially, a chemically modified oligonucleotide adapter (3’ adapter, 3A) is ligated to the 3’-hydroxyl group of the small RNA using a truncated T4 RNA ligase 2 (forming the product miRNA-3A). Following this, a second adapter (5’ adapter, 5A) is ligated to the 5’-phosphate group utilizing full-length T4 RNA ligase 1. The resulting biligated product (5A-miRNA-3A) serves as a template for reverse transcription into a single-stranded cDNA, which is subsequently amplified via polymerase chain reaction (PCR) to append flow-cell attachment sequences and indices for multiplexed sequencing.

The procedure outlined above does, however, come with the downside of introducing multiple sources of bias, particularly ligation bias. Ligation bias represents a pervasive challenge in small RNA sequencing ^6–8^. As small RNA library preparation relies on the direct ligation of adapters to a single, intact native fragment, sequence and secondary structure strongly influence ligation efficiency. As a consequence, final read counts and by extension the observed biological abundance of target species can be severely distorted. It is well established that the formation of intramolecular secondary structures within miRNAs and between miRNA/adapter pairs selectively enhances the capture of specific RNAs and diminishes others ^6,7,9^. Indeed, systematic evaluations show that alternative cDNA construction methods produce read counts differing by up to four orders of magnitude for the same input material ^10–12^. Ligation bias if unconstrained may unpredictably alter relative representational correctness depending on the composition of the input, as complex interactions between the diverse pool of miRNAs and adapter sequences dictate the overall reaction thermodynamics. Thus, even when analyzing highly related samples with similar composition, the final small RNA sequencing counts exhibit higher noise necessitating higher samples sizes for downstream analyses versus what would be the theoretical minimum required if miRNAs were converted to ligated products with no ligation bias present.

Efforts to mitigate ligation bias involve multiple distinct strategies developed over the past decade, including the use of degenerate bases, single-adapter circularization, and ligation-free methods. Protocols incorporating degenerate bases ^6,9,13–16^ introduce random nucleotides typically at the ligation boundary to effectively expand the pool of adapter sequences, increasing the probability of favorable interactions and efficient ligation for any given miRNA ^12^. Circularization approaches, such as that used in RealSeq AC ^17^ or others ^18^, involve ligating miRNAs with a single adapter followed by circularizing the products to bypass the traditional two-step ligation bottleneck. Other approaches utilize ligation-free template switching techniques ^19^ that use poly(A)-tailing and reverse transcriptase to bypass T4 ligase-mediated structural preferences entirely. However, these methods introduce their own set of reverse transcriptase and polyadenylation dependent biases while having lower sensitivity ^8,20–22^. It is still unclear what method is optimal to reduce ligation bias in practice, and the optimal one might depend on context and input sample complexity.

Standard commercial kits have a high background noise floor and encounter significant limitations when applied to low-input biological matrices such as liquid biopsies. In such samples, library preparation necessitates an increased number of PCR cycles. This combined with a lower ratio of desired products to adapters promotes the unwanted side effect of preferential formation and amplification of adapter-dimers, resulting in the potential loss of lowly abundant small RNAs from the final sequencing library ^12,23–25^. This strongly hampers reliable profiling in clinical and scarce-sample scenarios. Thus, there is a push towards protocols having higher library yields and sensitivities to ameliorate these issues and to improve library preparation robustness.

To resolve these bottlenecks, we engineered the MDX protocol, combining established bias-reduction approaches and practical workflow improvements with rationally designed, structure-forcing 5’ adapters that we identified to be key for improving overall ligation efficiency. Rather than solely relying on empirical screening, we utilized a bioinformatic analysis of the human miRNome to rationally design our 5’ adapters. By profiling the minimum free energy (MFE) distribution of natural miRNA-adapter interactions, we engineered 5’ adapters with controlled tethering designed to exceed a specific thermodynamic threshold. This targeted interaction physically overrides localized, unpredictable structural variations during the intermediate miRNA-3A state, forcing a universally favorable conformation that enables high-efficiency conversion of miRNAs into libraries even from low-input samples. Furthermore, it maintains mostly standard adapter sequences, allowing it to function as a drop-in replacement compatible with existing bioinformatic pipelines setup to work with Illumina sequencing kits.

## Results

### Bioinformatic modeling of intermediate ligation states guides rational 5’ adapter design

Our overarching goal was to minimize ligation bias while maintaining the high sensitivity required to enable robust miRNA marker detection from highly challenging, ultra-low-input biological matrices, such as cerebrospinal fluid (CSF) and cell culture supernatants (CCSN), or exosomes. To achieve this, we opted against testing individual improvements in isolation. Instead, we established a baseline protocol incorporating several principles known to be effective, aiming to build a synergistic workflow (Figure 1A). Our foundational method is conceptually based on a non-radioactive fluorescence-based PAR-CLIP (fPAR-CLIP) protocol ^26^ that in turn is derived from barcoded miRNA library construction protocol ^27^ based on the Illumina small RNA TruSeq adapter sequences. Specifically, this baseline physically separates the 3’ and 5’ ligation steps and utilizes a fluorescently-labeled 3’ adapter. Thus, adapter-dimer contamination, a critical issue that frequently plagues low-input library preparations and a frequent optimization target ^25,28,29^, can be minimized by preventing bulk unreacted 3’ adapter from entering the second ligation step but without the need to utilize radioactivity and a separate enzymatic digestion step to detect 3’ ligation products as in ^27^. Additionally, two degenerate bases at the ligation junctions are incorporated similar to ^13,14,26^. The use of degenerate junctional bases is a well-established approach known to dramatically reduce ligation bias ^13,14^, serving as an effective starting point for our optimization.

**Figure 1:**
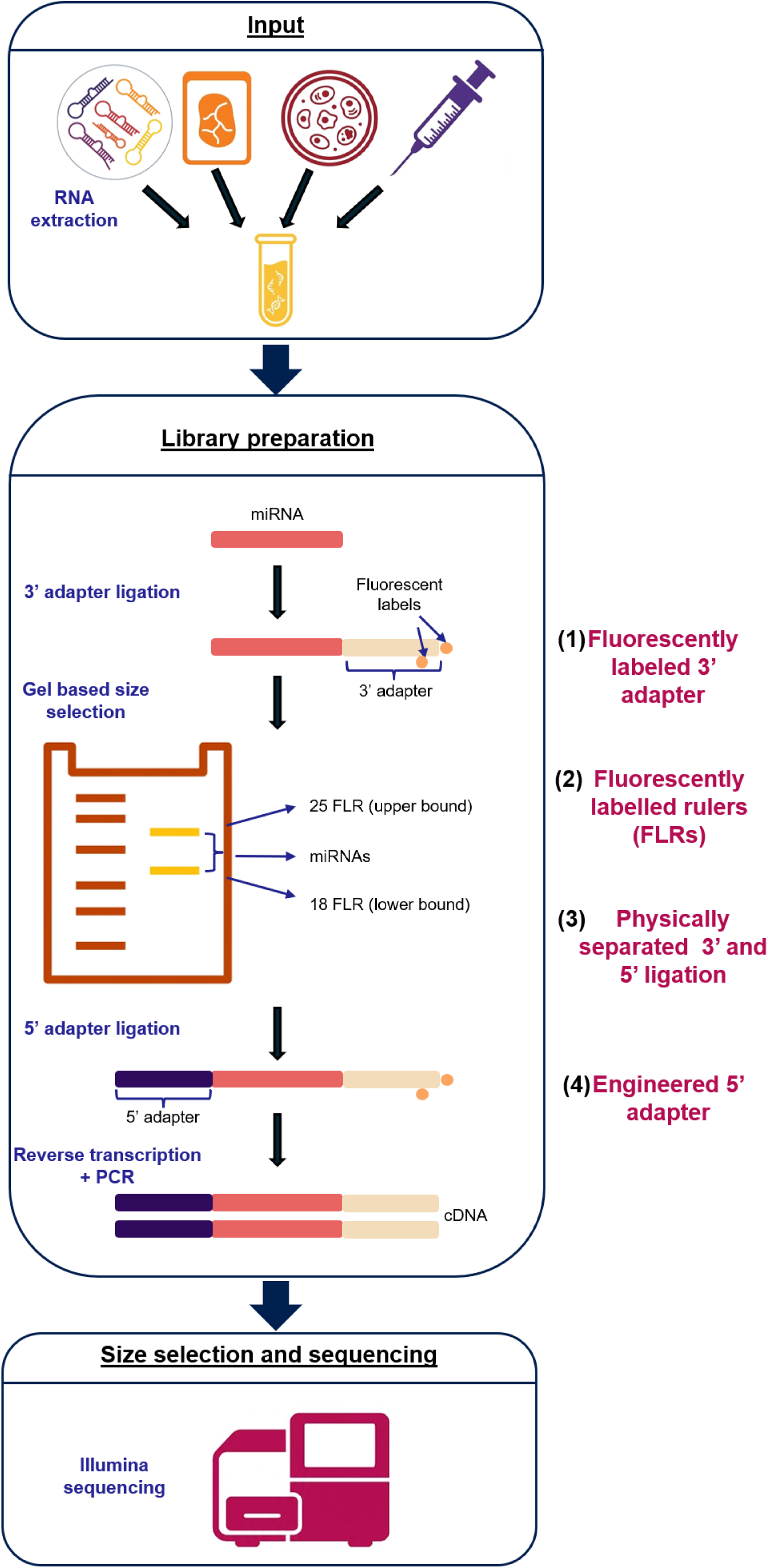
A schematic overview of the miRNA sequencing workflow, highlighting the key features included in the MDX protocol. A) Crucial features and innovations integrated in the MDX protocol. (1) Separating the 3′ and 5′ reactions decreases the formation of artifacts, such as primer dimers, while increasing library purity. (2) Incorporating a fluorescently labelled 3′ adapter enables direct visualization of intermediate ligation products with high inputs. (3) For low-input scenarios where fluorescently labeled intermediates are not visible to the naked eye, Fluorescent Ligation Rulers (FLRs) are added to 3′ ligated product prior to loading on denaturing PAGE gel. The two FLRs are compositionally similar the upper and lower end of the expected miRNA species, thus enabling more uniform product excision from the gel that takes lane-specific electrophoretic mobility and any artifacts into account. (4) Rationally designed 5′ adapters with thermodynamics-informed complementary sequences binding to the 3′ adapter in such a manner as to force a universally favorable cofold structure to form between the ligated 3′ adapter-miRNA and 5′ adapter to improve 5′ ligation yield and reduce bias.

During our early pilot experiments, we observed that the 3’ ligation step was inherently efficient (60+% conversion efficiency) and highly malleable to further improvement through the optimization of reaction conditions. On the other hand, the 5’ ligation reaction proved much less efficient, with conversion yields frequently falling below 30%, even as low as 10% under certain conditions. Such low numbers also made us suspect that ligation bias might be severe as well despite the presence of degenerate junctional bases in our baseline adapter sequences. We reasoned that either lack of collisions and/or unfavorable secondary structures between the post 3’ ligation miRNA product (miRNA-3A) and the 5’ adapter were driving this inefficiency. This thinking is consistent with previous structural studies showing that types of secondary structures between miRNAs and adapters can not only influence ligation efficiency but also bias^9,15^.

Building upon this, we hypothesized that controlled tethering could not only bring the interacting molecules together but also override localized, unpredictable structural variations in the miRNA-3A intermediate. This would address both fronts simultaneously: artificially increasing effective local concentration to boost absolute yield, while flattening the thermodynamic landscape to minimize bias. Based on this idea, we engineered improved 5’ adapters featuring rationally designed regions complementary to 3’ adapter and evaluated their ligation efficiency and bias. To test our hypothesis, we designed two distinct complementary domains within the 5’ adapter, designated ’A’ and ’U’, that target the second half and 3’ ends of the 3’ adapter, respectively, adding one or both to the 5’ or 3’ end of the core 5’ adapter sequence (Figure 2A). We omitted targeting the 5’ end of the adapter due to presence of an optional barcode for sample pooling ^6^ and the fact that follow-up 5’ sequence of the native Illumina 3’ adapter was GC poor (TGGAATT) that would have necessitated an impractically long complementary sequence. The sequences of these domains, while naturally constrained by the available sequence complexity of the 3’ adapter, were not selected at random. Instead, we used a thermodynamics-informed bioinformatic approach to guide necessary length/GC composition of the engineered domain to ensure the interactions were strong enough.

**Figure 2:**
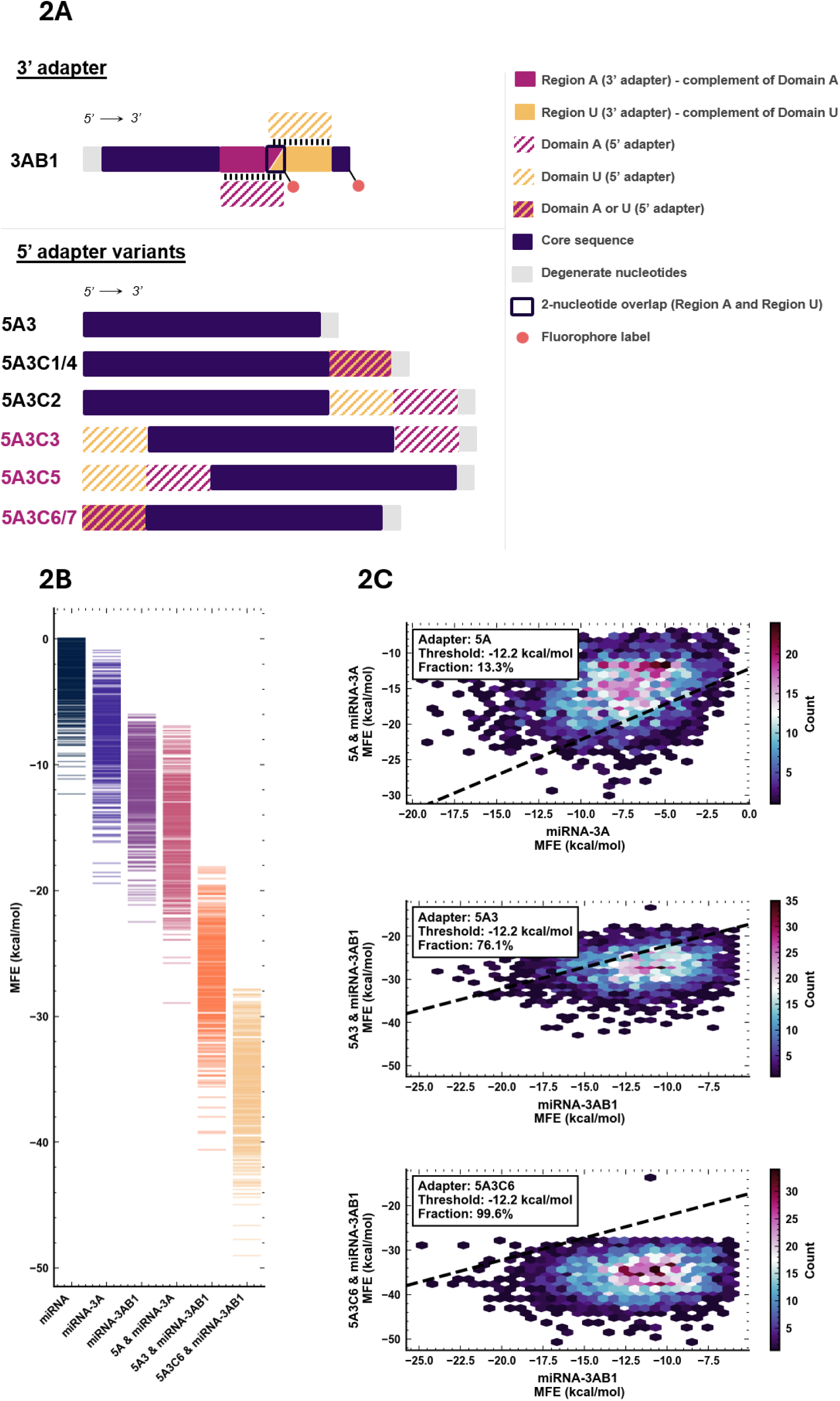
Structural design and minimum free energy (MFE) modeling of engineered 5′ adapters. A) Schematic of engineered 5′ adapter variants targeting the 3′ adapter (3AB1) via complementarity domains to maximize ligation yield and minimize ligation bias. Both adapter types contain two degenerate junctional bases (NN) to further reduce bias, while core sequences maintain Illumina platform compatibility. Except for the baseline control (5A3), all variants (5A3C1 - 5A3C6) feature non-overlapping Domains A and U. Their corresponding targets on 3AB1 region A (2^nd^ half of the adapter) and region U (3′ end) - overlap by exactly 2 nucleotides. This overlap carries no biological function and is a purely structural constraint of the 5′ primary sequence design. Black lines indicate direct sequence complementarity. Variants are ordered top-to-bottom, starting with control 5A3, followed by two groups distinguished by domain localization at the 3′ or 5′ terminus. Deep magenta annotations highlight the empirically determined to be superior 5′-terminal domain placement. B) Predicted minimum free energy (MFE) distributions (500 miRNAs subsampled for visual clarity) determined via ViennaRNA for the self-folding and co-folding structures of all annotated human miRNAs in miRBase v22.1 and their intermediate post-3’-ligation (miRNA-3A) products. C) Joint MFE distributions of the above and calculated high-conversion (95%) fractions for the standard 5A Illumina adapter (top), the baseline 5A3 adapter (middle), and the engineered 5A3C6 adapter (bottom) as an example of one of the custom engineered adapters relative to the total miRNA-3A intermediate pool. The horizontal dashed line defines the custom metric, thermodynamic threshold, MFE_gap_, of -12.2 kcal/mol computed to conservatively achieve 95% conversion into the heterodimeric complex at equilibrium under 5’ ligation conditions.

First, using ViennaRNA ^30^, we evaluated the predicted minimum free energy (MFE) distribution for the self-folding and cofolding structures of all human miRNAs annotated in miRbase v22.1 (2656 in total) and their intermediate miRNA-3A products (Figure 2B). Second, we plotted the joint MFE distributions between the 5A adapter and miRNA-3A intermediates (Figure 2C). To define a conservative thermodynamic threshold where the engineered adapter effectively outcompetes native folding, we modeled the threshold, labeled as MFE_gap_ (see Methods section for more details), required to achieve a theoretical 95% conversion into the heterodimeric 5A & miRNA-3A cofold complex (“&” denotes cofolding) at equilibrium. This established a critical threshold of -12.2 kcal/mol for conditions encountered during 5’ ligation (Figure 2C, dashed line). By engineering the 5’ adapter domains such that the modified adapters meet or exceed this binding strength for almost every miRNA considered (Figure 2C, bottom), we conservatively ensure they overpower any native, competitive intramolecular folding of the miRNA-3A intermediates. This modeling demonstrates that the core Illumina small RNA TruSeq sequences fail to clear this threshold for most miRNAs, with the core 5A adapter achieving a high-conversion fraction of just 13.3% (Figure 2C, top). While our baseline protocol (5A3 + 3AB1, with degenerate bases and 5’ barcode extension to the 3’ adapter) significantly shifts this distribution to 76.1% (Figure 2C, middle), it remains imperfect.

Crucially, because our core objective was to flatten the thermodynamic landscape and override miRNA sequence-dependent folding biases, we conservatively designed our engineered domains to be the overwhelmingly favored structures while only considering intermolecular interactions. This slightly underestimates the driving force for complex formation and builds in a buffer for situations where competing interactions that would act as barriers (i.e., heterodimers with other oligonucleotides that cannot be reliably modeled) need to be broken first. While this approach serves as a stringent proxy engineering metric rather than an exhaustive ensemble partition function simulation, it established a highly reliable design criterion for successful engineering of the ’A’ and ’U’ domains.

### Spatial arrangement of complementarity domains dictates ligation performance

We structured the initial screening to determine if the diverse secondary structures formed by the miRNA-3A ligated complexes necessitated different tethering points for optimal ligation and what optimal placements are for our designed domains in 5’ adapter (Figure 2A). These were either placed alone/together at the 3’ or 5’ end of 5’ adapter in some designs or split such that each position contained one domain each.

This last approach aimed to evaluate whether tethering flexibility was necessary to achieve optimal ligation performance. Together we evaluated 7 variants, labeled 5A3C1 through 5A3C7.

The screening process revealed that both ligation efficiency and ligation bias are highly sensitive to the spatial arrangement of these domains (Figure 3), but the magnitude differs. Specifically, ligation efficiency (that is the overall input conversion rate) was dramatically improved with the addition of almost any engineered domain, regardless of its type, count, or position except for 5A3C4 (Figure 3A). Densitometric measurements revealed only small differences among the well-performing modified variants, with conversion yields reaching the 76-83% range compared to just 44% for the baseline 5A3 adapter.

**Figure 3:**
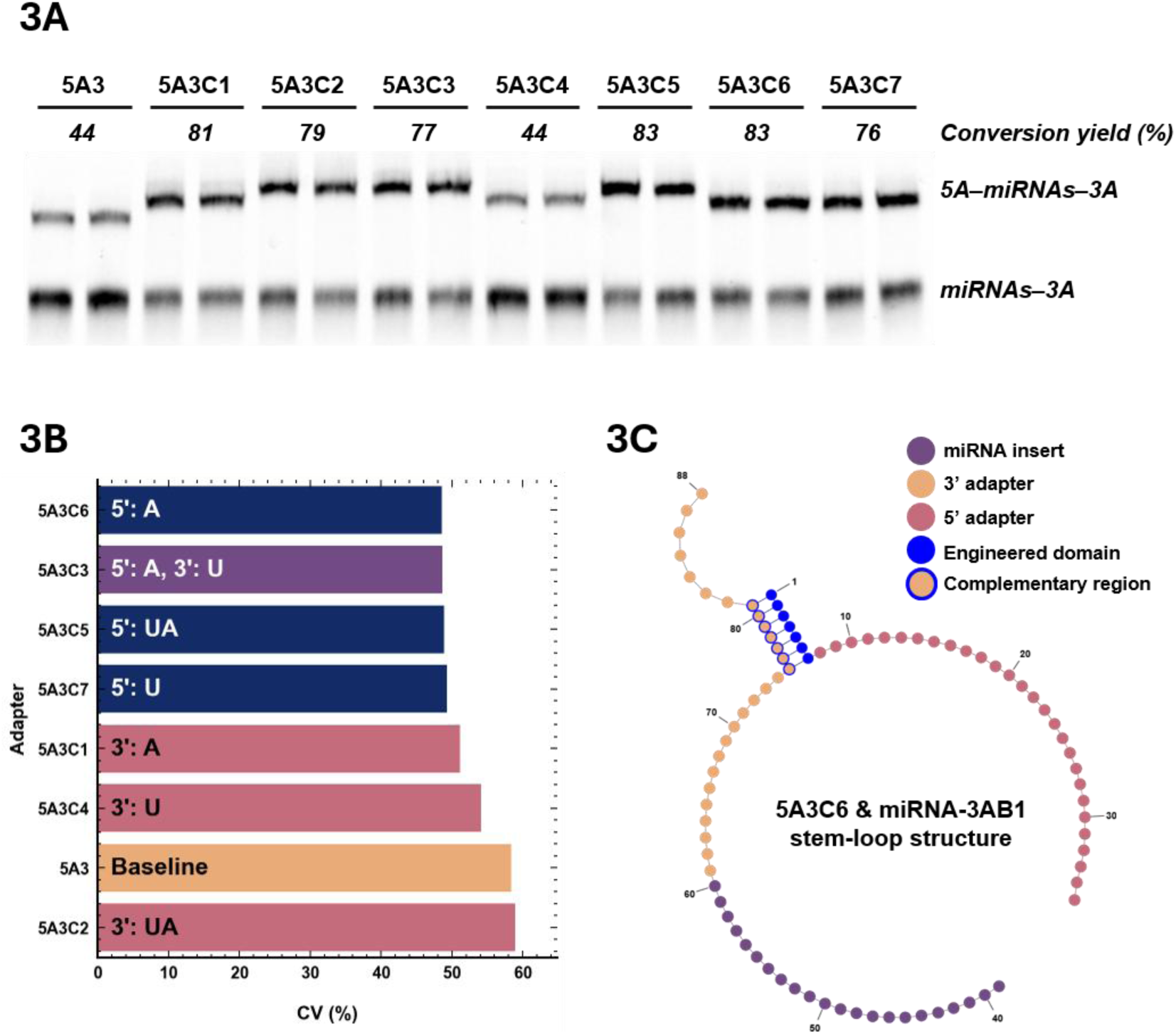
Spatial arrangement of complementary domains dictates ligation bias. A) Representative denaturing polyacrylamide gel electrophoresis analysis demonstrating the 5’ ligation conversion efficiency of the unmodified 5A3 baseline adapter versus custom modified variants (5A3C1 through 5A3C7) reacting with the intermediate miRNA-3A pool to generate the final biligated 5A-miRNA-3A product. Numbers underneath group labels represent the calculated conversion yield (average of 2 technical replicates). Conversion yield calculation was adjusted to consider that the miRNA-3A intermediate was in 67% excess vs. 5’ adapter concentration in this particular experiment. B) Evaluation of representational ligation bias (average of 2 technical replicates) across the baseline control (5A3) and 7 modified adapter designs (5A3C1 through 5A3C7), quantified as the Coefficient of Variation (CV, %) of the ratio between observed and expected read counts using a synthetic miRNA reference panel. Labels indicate whether the engineered complementarity domains (A, U, or UA) were appended to the 5’ or 3’ terminus of the core adapter sequence. C) Predicted secondary structure of the prototypical 5A3C6 adapter and miRNA-3AB1 intermediate considering only the designed complementarity domain. The illustration highlights how adding a 5’-terminal complementarity domain is predicted to generate a split-loop stem-loop conformation that physically draws the 5’ and 3’ ligation junctions together through distal tethering involving 5’ end of the 5’ adapter and 3’ end of the 3’ adapter.

Contrarily, analyzing ligation bias, measured as the ratio between the observed versus expected number of reads based on the number of unique miRNAs in the synthetic mix used as input scaled to the effective sequencing depth, revealed a highly specific, position-dependent pattern of results (Figure 3B; Supplementary Figure 1A). Note that while many prior works utilized the miRXplore Universal Reference panel (Miltenyi Biotec, US) for ligation bias studies, it is no longer commercially available and thus we had to develop an internal alternative that was suitable for evaluating differences between tested adapter designs (88x or 150x miRNA panel depending on experiment, described in more detail later; for composition see Supplementary Table 1).

While all engineered 5’ adapters other than 5A3C2 achieved a lower ligation bias than the baseline 5A3 adapter (CV = 58.4%), performance varied based on domain positioning. Adapters with one or two domains added strictly to the 3’ end performed noticeably worse than those where at least one domain was added to the 5’ end.

Secondarily, designs incorporating the ‘U’ domain underperformed compared to those with the ‘A’ domain, a trend clearly visible in 3’ modified adapters. However, in the case of designs with 5’ modifications, these performance differences were inconsequential, within 1%. Adding an engineered domain to the 5’ end of the 5’ adapter thus acted as an ’equalizer’ in terms of performance, rendering this approach unambiguously the best choice; adding a second domain anywhere else offered no additional benefit. This behavior is strongly supported by clustering analysis (Supplementary Figure 1B), that shows that modified adapters with a domain added to the 5’ end cluster tightly together within their per-miRNA ligation profiles. Consequently, we selected 5A3C6 (CV = 48.6%) as the prototypical adapter for our final design not only because of its overall performance but also due to its structural simplicity.

The specific architecture of 5’ modified adapters exemplified by 5A3C6 is predicted to form a split-loop stem-loop structure intermediate, physically drawing the 5’ and 3’ ligation junctions together through what we call “distal tethering” (Figure 3C). Because the engineered complementary sequence of 5A3C6 is located upstream of the PCR primer binding site, this added sequence is not incorporated into the final PCR amplicon. This architecture has the added benefit of not increasing the final library fragment size, thereby avoiding any potential downstream purification complications such as reduced elution efficiency or worsened electrophoretic separation between adapter dimers and insert-containing amplicons.

### Engineered adapters change thermodynamic and structural drivers of ligation bias

In a separate sequencing experiment (Supplementary Figure S3), we analyzed correlations between structural features – including the cofold type classification used by ^15^ to classify oligonucleotide interactions in the context of small RNA library preparation – and selected adapter sequences, looking at both the interactions of baseline unmodified Illumina TruSeq adapters and chosen custom adapter variants (5A3, 5A3C3, 5A3C6). To this effect, we utilized ViennaRNA’s cofold function and a custom Python script that converts dot-bracket notations into cofold types. To improve accuracy, degenerate bases in adapter sequences were replaced with empirically most dominant variants encountered in sequencing data itself on a per-miRNA basis. We evaluated Spearman correlations between empirical bias and miRNA self-folding, all pairs of bimolecular cofolding, length of loop with the ligation junction for the predicted 5A & miRNA-3A cofolds, miRNA GC content, and miRNA length (Supplementary Figure S3A).

For the original Illumina adapter sequences (5A TruSeq), a highly significant negative Spearman correlation was observed between ΔMFE (the driving force of complex formation according to our model) and bias (ρ = -0.55, p = 3.36e-8), while a weaker correlation was found for the unadjusted cofold MFE value of 5A & miRNA-3A (ρ = -0.46) (Supplementary Figure S3B). This suggests that our approach has higher explanatory power for predicting what miRNA species will be overrepresented with libraries preparation methods with high levels of ligation bias. Additionally, a moderate positive correlation existed with the native miRNA MFE (ρ = 0.32), that is unstructured miRNAs tended to be overrepresented in final libraries. Conversely, for our custom adapters, these structural relationships collapsed or reversed, likely due to the much lower levels of bias observed for them. The Spearman correlation for ΔMFE increased above zero for the baseline 5A3 adapter (ρ = 0.10, p = 0.350) and the prototypical 5A3C6 adapter (ρ = 0.18, p = 0.100), before significantly reversing for the 5A3C3 adapter (ρ = 0.26, p = 0.018), a variant that is predicted to form even stronger secondary structures on average. The correlation with the 5A & miRNA-3A cofold MFE similarly shifted to positive values for 5A3C6 (ρ = 0.19) and 5A3C3 (ρ = 0.28). A strong, highly significant positive correlation with native miRNA GC content was also uniquely observed for the baseline 5A3 adapter (ρ = 0.51, p = 8.16e-6). We also investigated whether loop length of the predicted junction loop or bulge of 5A & miRNA-3A affected bias, but no significant correlations could be found for any of the adapter investigated.

Finally, we examined whether specific cofold types dictated representational bias (Supplementary Figure S3C-D) as was previously reported ^9,15^. Cofold type distribution (Supplementary Figure S3C) did differ between baseline Illumina TruSeq adapters and our custom ones suggesting cofold types might explain the differences in empirically measured bias (Supplementary Figure S3D). However, for no adapter combination investigated did Mann-Whitney test show statistical significance for such an effect.

Thus, our data highlights that cofold type by itself is not the only determinant of ligation bias and strength of interaction between species is a strong, independent predictor of miRNA library representation at least in libraries with high level of observed bias.

### Engineered adapters are compatible with Unique Molecular Identifiers (UMIs)

Additionally, we evaluated whether our prototypical 5A3C6 design was in principle compatible with including a UMI sequence that can be used to correct for reverse transcription and PCR amplification-induced bias ^31^. To this end, we evaluated a 5A3C6 variant we dubbed 5A3C6U that supplants our engineered complementary sequence with an anchor sequence and a segmented (to reduce aggregation propensity) 10 nt-long UMI sequence near 3’ end of the 5A right before the junction (Supplementary Figure S2A). This modified adapter shows bias near identical to the parent adapter 5A3C6 (Supplementary Figure S2B-C) at both high and low levels of input. Importantly, the distribution of individual nucleotides in the UMI region is on average broadly random (Supplementary Figure S2D) and even miRNA-specific analysis (evaluated as bit entropy per position per miRNA) does not show appreciable deviations from randomness (Supplementary Figure S2E). While a few specific sequences show evidence of modest bias (bit entropy delta of 0.2-0.4 out of 2.0 maximum), these do not appear to affect miRNA representation in final libraries as they are present uniformly across all miRNAs irrespective of their abundance rank in the final library. Thus, if UMIs are required, for example for ultra-low input processing scenarios, our design permits their incorporation and use. However, since it has been shown previously that RT-PCR bias is generally not of a significant concern for general miRNA library preparation ^6,32,33^, and no commercial kit we used for benchmarking (see later sections) uses UMIs, we opted to use the simpler 5A3C6 adapter for all further experiments, guaranteeing strict parity in downstream bioinformatic processing.

### Optimization of two-stage reaction and purification conditions increases biligated product recovery

While the 5A3C6 adapter successfully resolved the biochemical bias of the 5’ ligation step, applying this innovation to ultra-low-concentration liquid biopsies, such as CSF, required a highly robust physical workflow. As mentioned before, the core design principle of our protocol is the physical separation between the 3’ and 5’ ligation steps ^6^. This separation is critical to minimize adapter dimer contamination, a pervasive issue that typically plagues low-input small RNA library preparations. However, this separation strategy inherently necessitates an intermediate PAGE gel purification step, which comes at the cost of reduced throughput and significant potential for material loss and increased technical variance (multiple handling steps, gel elution, precipitation, etc.). Thus, we set out to optimize these steps to improve ligated product generation and retention. Note that these experiments were performed in parallel with structural investigations and therefore they utilize 5A3/3AB1 adapter pair and not 5A3C6/3AB1 as was later identified to be optimal.

Initial investigations utilizing standard two-stage ligation conditions revealed conversion yields ranging from 10% to 75% across the primary stages, with gel elution and 5’ ligation being the main sources of material loss (Figure 4). Consequently, only low single digit percent of the initial input material was successfully converted into the 5’/3’-biligated product necessary for downstream library amplification. To overcome this severe loss of final product, we systematically optimized both the ligation reactions and the gel elution procedure. Surprisingly, we found that very few modifications had beneficial effects. The most impactful changes were balancing PEG and DMSO concentrations during 3’ ligation (15% PEG, 20% DMSO found optimal) (Figure 4A), adding 3 freeze-thaw cycles before initiating ligated product elution from gel pieces between 3’ and 5’ ligation steps (Figure 4B), and prolonging 5’ ligation from 1 hour to 4 hours (Figure 4C). Collectively, these targeted optimizations when combined can result in a hypothetical median ∼4.8-fold increase (from 2.7-9.9% to 21.2-32.8%; Figure 4D) in the final yield of 5’-3’ biligated products, substantially improving the overall efficiency of the library preparation.

**Figure 4:**
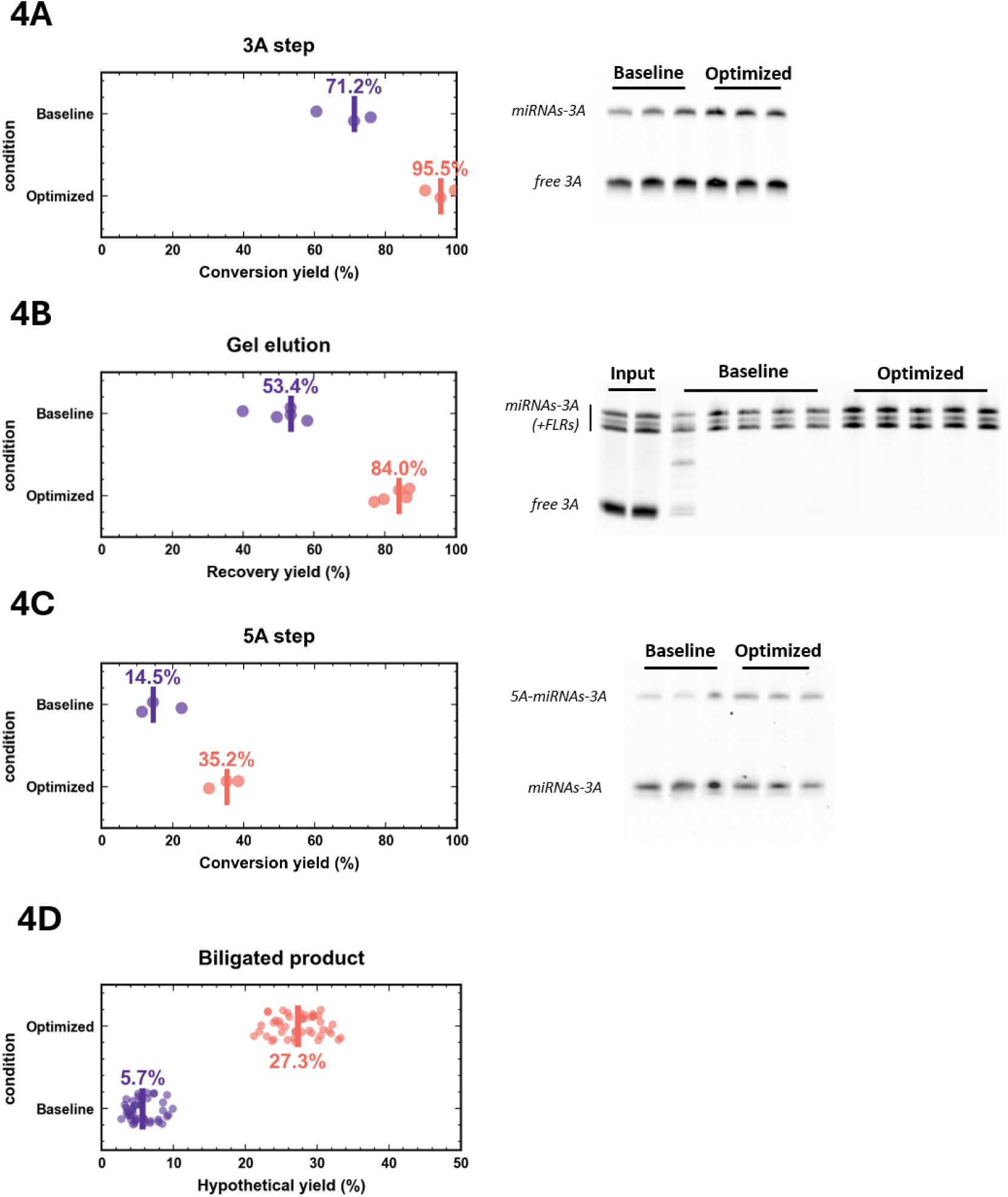
Systematic optimization of two-stage reaction and purification conditions increases biligated product recovery. A) Ligation conversion yield (%, 3 technical replicates) and representative polyacrylamide gel image comparing the efficiency of the 3’ ligation step under baseline reaction (15% PEG only) parameters versus optimized conditions (15% PEG, 20% DMSO). Conversion yield calculation was adjusted to consider that the 3’ adapter was in 100% excess vs. miRNA concentration in this particular experiment. B) Recovery yield (%, 5 technical replicates) and representative gel image evaluating intermediate target retention during gel purification, comparing standard elution procedures against an optimized protocol incorporating three targeted freeze-thaw cycles prior to product elution. Both ligation and FLR intermediates were used to compute the final yield. C) Conversion yield (%, 3 technical replicates) and representative gel image of the 5’ ligation reaction comparing standard 1-hour baseline incubation time against an optimized, prolonged 4-hour incubation time. D) Combined dot plot illustrating the total hypothetical yield (%) of the final 5’-3’ biligated product achieved by computing all possible measured baseline values per evaluated step versus fully optimized workflow steps.

### Workflow robustness is improved by utilizing rationally engineered fluorescent ligation rulers

Despite these improvements in yield, the intermediate gel excision step remained a potential source of technical variation between library preparations. Traditional methods for band excision rely either on hazardous radiolabeled oligonucleotides ^27^ or on imprecise visual estimation based on a molecular weight ladder, both of which limit throughput and reproducibility. While our adapters are fluorescently labeled, in most scenarios we were internally targeting (library preparation from low input liquid biopsy samples), the amount of miRNA-3A intermediate is too low to visualize unlike in ^26^, even with prolonged imaging times. To rectify this, we developed rationally designed "Fluorescent Ligation Rulers" (FLRs). These in-lane fluorescent oligos, fluorophore-matched with our 3’ adapters, correspond to the precise upper and lower boundaries of the target ligated miRNA species (25 and 18 nt, respectively) and are "pre-ligated" in sequence and labeled identically to the target products to ensure identical electrophoretic migration (Figure 5A). When added to individual wells prior to electrophoresis, FLRs provide in-lane molecular weight markers that correct for any gel-specific migration artifacts or deformities. As FLRs can be labeled with fluorophores that can be excited with green LED light (i.e, AF488), with proper instrumentation and safety glasses, an operator can observe the bands directly while performing excision (Figure 5B), greatly improving speed and robustness of this crucial but otherwise hard-to-standardize step.

**Figure 5:**
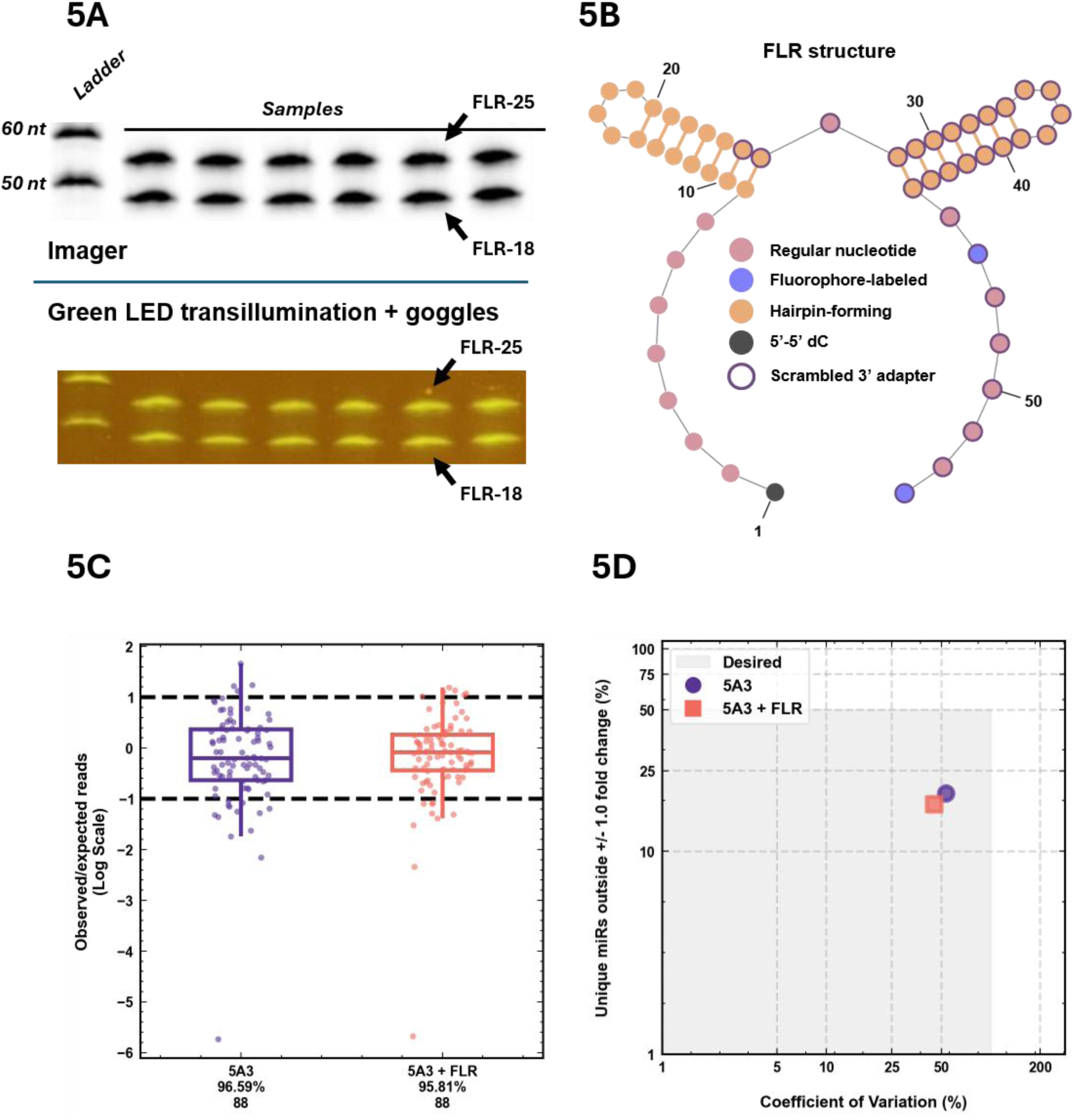
Structural features and workflow validation of rationally engineered Fluorescent Ligation Rulers (FLRs) A) Upper part shows a fluorescence image of in-lane Fluorescent Ligation Rulers (FLRs) added directly to individual wells prior to purification, serving as co-migrating molecular markers that delineate the precise upper (25 nt) and lower (18 nt) migration boundaries of target small RNA species. Bottom part shows the same polyacryalamide gel as seen by the operator (captured by a regular phone camera) under green LED transillumination as seen while wearing safety goggles. B) Secondary structure model of the engineered FLRs, highlighting the scrambled 3’ adapter core, dual hairpin-forming domains designed to lock the sequence under ligation conditions, and the 5’-terminal 5’-5’ dC linkage to ensure downstream enzymatic inertness. C) Boxplots depicting ligation bias of libraries prepared without (5A3) or with FLRs (5A3 + FLR), expressed as the log2-transformed ratio of observed to expected read counts (1 technical replicate). First row of numbers underneath group labels indicate the mapping rate. Second row of numbers underneath group labels indicate number of unique miRNAs detected for individual groups. Horizontal dashed lines define the +/- 1 log2 fold change low-bias range. D) Scatter plot of above summarizing protocol performance by tracking the percentage of unique miRNAs falling outside a +/- 1 log2 fold change range against the overall Coefficient of Variation (CV, %), demonstrating that the addition of FLRs (5A3 + FLR) introduces no new sources of bias compared to libraries prepared without them (5A3).

Because FLRs are co-eluted with the 3’-ligated miRNAs, a crucial design requirement was ensuring they remain completely inert during downstream 5’ ligation, reverse transcription, and PCR steps, where their total amount would otherwise vastly outnumber the true ligation products. To achieve this inertness, we rationally designed the FLR sequences with multiple properties (Figure 5C). First, we scrambled the 3’ adapter sequence within the FLRs to weaken any predicted secondary structures with 3’ adapter and 5’ adapter and engineered the sequence with two separate hairpin-forming regions. This split-hairpin design ensures that the FLRs completely melt under denaturing Urea-PAGE conditions to act as precise molecular rulers yet are predicted to form specific secondary structures under standard ligation conditions that lock more than 80% of the sequence from interacting with other oligonucleotides (MFE = -16 kcal/mol), preventing their interference in subsequent enzymatic steps. Furthermore, the 5’ terminal nucleotide of the FLRs features an unnatural 5’-5’ dC linkage, rendering them functionally unusable as substrates for 5’ ligation but having electrophoretic mobility comparable to native miRNA-3A intermediates of the same size.

Crucially, comparative testing of libraries prepared from a synthetic miRNA panel with and without the use of FLRs demonstrated no detrimental effects of FLRs on final library composition (Figure 5D-F). Mapping rates, ligation bias, and library quality were all comparable between libraries prepared with or without FLRs in the workflow. In addition, bioinformatic analysis of the final sequencing data confirmed complete absence of FLR-derived fragments (0 FLR fragment read counts out of ∼3 million reads in total) in the target libraries, validating their engineered inertness during downstream amplification.

### MDX protocol minimizes ligation bias in comprehensive synthetic benchmarks

With the optimized workflow (FLRs, optimized reaction/elution conditions) and adapter selection (the fluorescent 3AB1 + 5A3C6 engineered adapters) combined into what we call the MDX protocol, we wanted to perform an in-depth benchmarking study against leading commercially available small RNA sequencing kits. To evaluate true library preparation bias, we developed an in-house synthetic panel of 150 human miRNAs (Supplementary Table 1) curated from miRBase version 22.1^34^. This equimolar panel was bioinformatically designed to represent the structural diversity of the human miRNome, including sequences with extreme GC content, varied lengths, strong self-folding energies, and pairs predicted to form strong heterodimers (Supplementary Figure S4A-D). Using this 150x ground-truth panel, we benchmarked our MDX protocol against three commercially available and widely used kits: Illumina Small RNA TruSeq, Revvity NEXTflex v4, and RealSeq AC. All are Illumina-based in terms of core adapter sequences and thus analogous to our custom approach; the latter two additionally include modifications that improve ligation efficiency and bias compared to the Illumina baseline ^17,35^. Evaluations were performed using 10 ng and 1000 ng total RNA equivalent inputs (see Methods for details), reflecting the manufacturer-recommended ranges for these commercial kits. To ensure a fair and direct comparison of ligation efficiencies and yields, all protocols were unified post-reverse transcription, utilizing identical PCR amplification steps, enzymes, cycle numbers, and downstream PAGE purification procedures as these can be in principle freely swapped between protocols.

The MDX protocol, with both 1000 ng and 10 ng total RNA equivalent input, significantly outperformed the commercial alternatives, exhibiting the lowest overall ligation bias (Figure 6A), the lowest CV for miRNA bias as well as the fewest percentage of miRNAs with highly biased representations (Figure 6B). With the MDX protocol, ∼75% of the miRNAs in the synthetic panel fell within a +/- 1 log2 fold change range, indicating that most miRNA species were overrepresented by no more than 100% or underrepresented by no more than 50%. Among the commercial options, RealSeq demonstrated the lowest bias with ∼50% of miRNAs in the same desired bias range, whereas both the NEXTflex and TruSeq kits produced highly skewed read distributions with most miRNAs significantly underrepresented in their final libraries. Analysis of inter-input amount correlations (Pearson, log-log transformed reads per million mapped, RPM, values averaged from technical replicates) from the same experiment (Figure 6C) revealed high positive correlation (r = 0.95). This was held true for the commercial protocols as well (r = 0.91-0.98) confirming that representational bias did not shift for any protocol by reducing the input by a factor of 100. On the other hand, inter-protocol correlations were much lower (r < ∼0.8) and especially RealSeq (Figure 6C) had generally low correlations (r < 0.55) with all other protocols likely reflecting its different ligation approach through circularization fundamentally affecting the bias profile of the prepared libraries.

**Figure 6:**
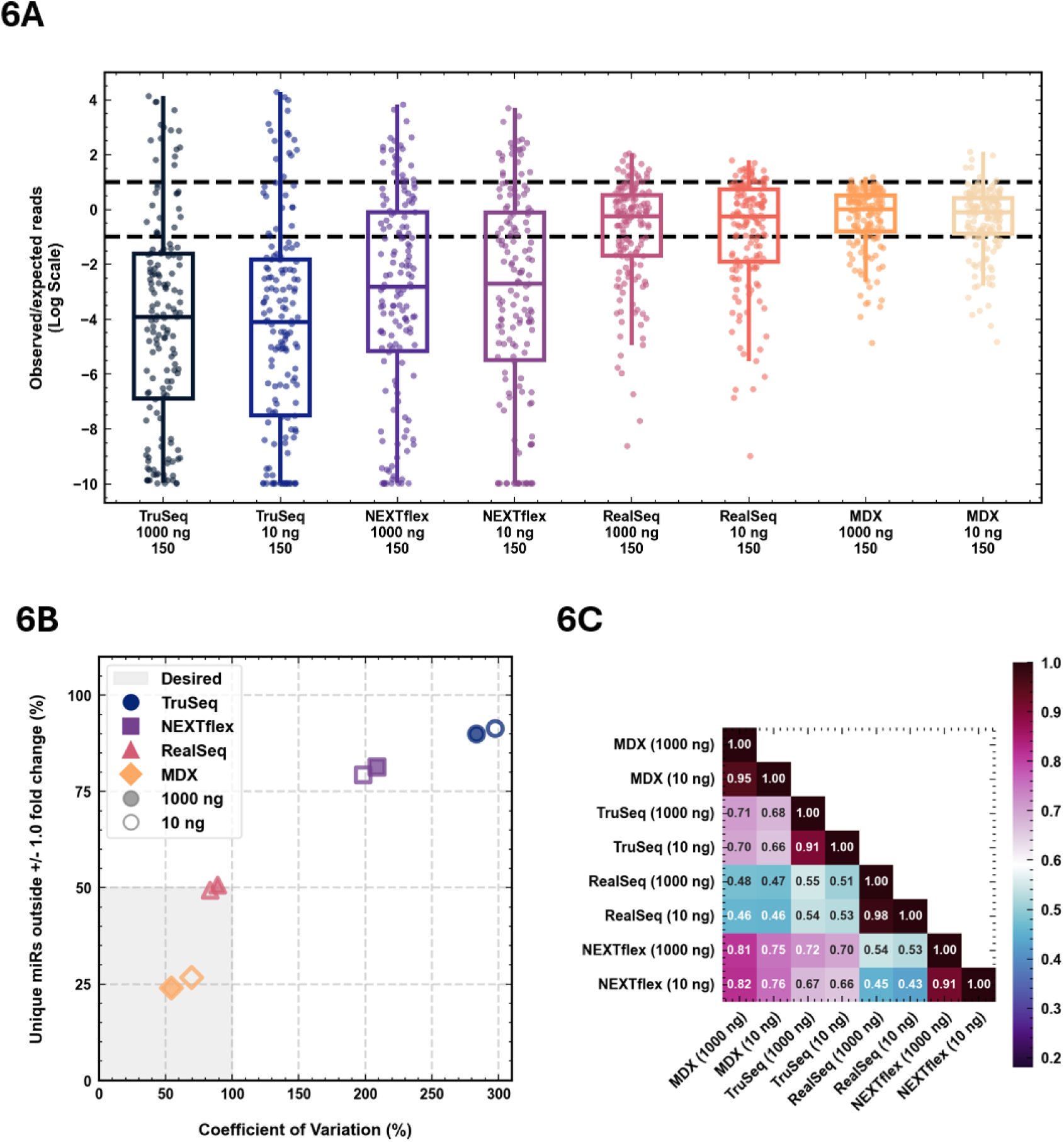
Comparative performance evaluation of the MDX protocol and commercial small RNA sequencing kits on a synthetic human miRNA panel. A) Boxplots depicting ligation bias (average of 2 technical replicates), expressed as the log2-transformed ratio of observed to expected read counts, for TruSeq, NEXTflex, RealSeq, and MDX library preparation methods across 1000 ng and 10 ng total RNA equivalent inputs using a 150x equimolar synthetic panel. First row of numbers underneath group labels indicate the input amount. Second row of numbers underneath group labels indicate number of unique miRNAs detected for individual groups. Horizontal dashed lines define the +/- 1 log2 fold change low-bias range. B) Scatter plot of above summarizing benchmarking performance by plotting the percentage of unique miRNAs residing outside a +/- 1 log2 fold change window against the overall Coefficient of Variation (CV, %) across tested protocols and input amounts. C) Heatmap from the same experiment displaying Pearson correlation coefficients (r, log-log transformed RPM values) calculated across independent protocols and input concentrations to evaluate library consistency and protocol-specific bias differences.

As our protocol is more complex than the commercial alternatives used in our benchmarking, reproducibility is of key concern, especially when working with smaller, challenging sample sets where bias reduction might be of the greatest benefit. In additional experiments focused on technical reproducibility and sensitivity, we evaluated the detection limits of our protocol and inter-replicate correlation. We successfully generated libraries from as little as 0.1 ng of total RNA equivalent input (Supplementary Figure S5A); however, ligation bias worsened below 0.5 ng input (Supplementary Figure S5B) even though replicates at any given input concentration remained highly correlated (Supplementary Figure S5C). This suggests that the higher variance is not due to operator error with low input but instead reflects a shift in ligation dynamics. Indeed, when explicitly testing inter-replicate variance with 6 technical replicates evaluated at 1 ng total RNA equivalent (Supplementary Figure S5D), the correlation was high (r = 0.91-0.99) confirming the high robustness of the assay.

### MDX protocol enhances biomarker detection in complex, low-input matrices

Finally, to evaluate protocol performance in low-input and complex biological matrices, libraries were prepared from cerebrospinal fluid (CSF; 400 µl), cell culture supernatants (CCSN; 1000 µl) of cerebral organoid cultures, and formalin-fixed paraffin-embedded (FFPE; 100 ng) breast cancer tissues. Samples used were pools of multiple individual specimen samples to guarantee identical starting input composition for all benchmarked protocols and to approximate the average performance on any given matrix. The MDX protocol yielded uniformly high miRNA mapping rates and total identified miRNAs across all matrices (Figure 7). Compared to the other benchmarked protocols, the greatest difference in mapping rates was observed in case of CSF as input - MDX protocol achieved a miRNA mapping rate of almost 70%, whereas the commercial protocols yielded mapping rates between 0.6% and 7.5% (Figure 7A).

**Figure 7:**
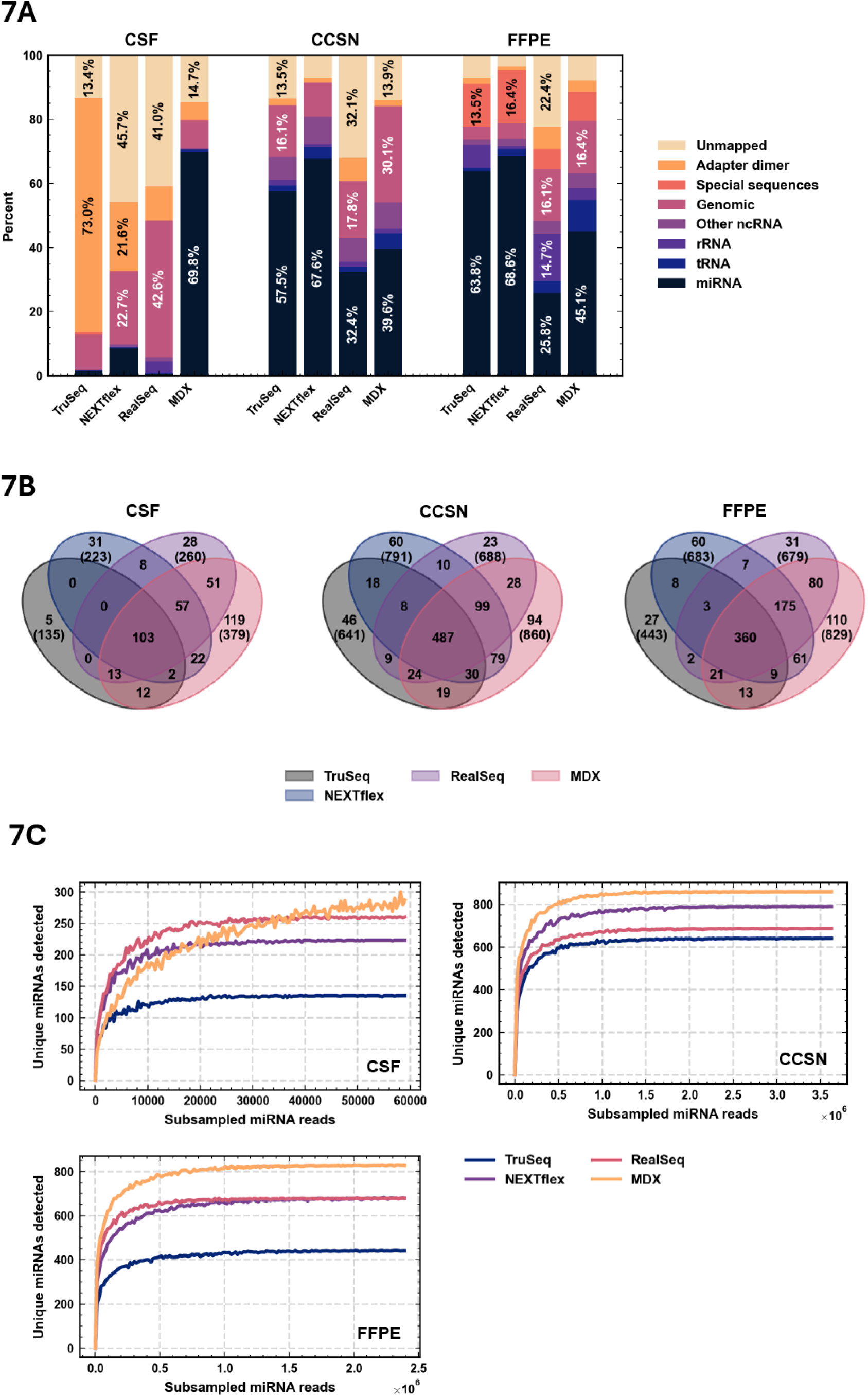
Comparative performance evaluation of the MDX protocol and commercial small RNA sequencing kits on challenging biological matrices. A) Stacked bar charts illustrating the relative percentage distribution of sequencing reads mapping to specific RNA biotypes, genomic elements, and technical artifacts across CSF, CCSN, and FFPE sample types (1 technical replicate each) using TruSeq, NEXTflex, RealSeq, and MDX library preparation methods. Numbers inside individual bar segments indicate the exact percentage for any annotation category capturing greater than 10% of the total sequencing reads. B) Four-way Venn diagrams showing the unique and overlapping miRNA features successfully detected by each method within the CSF, CCSN, and FFPE sample pools. The standard outer petal label contains the number of unique miRNAs exclusive to that specific protocol, while the value underneath in parentheses denotes the total number of unique miRNAs identified by that protocol (representing the mathematical sum of all overlapping petals involving that particular protocol). C) *In silico* downsampling analysis to evaluate unique miRNA detection sensitivity independent of variations in effective sequencing depth across CSF, CCSN, and FFPE sample types. The plots show the number of unique miRNAs recovered across resampled miRNA reads in 2000 count steps, demonstrating the inherent analytical sensitivity of the TruSeq, NEXTflex, RealSeq, and MDX library preparation methods at uniform sequencing depths.

Furthermore, the higher library quality produced by MDX protocol is also exemplified by the presence of only ∼5% adapter dimer reads. On the opposite end of the spectrum, the TruSeq protocol generated >70% adapter dimer reads from identical CSF input. The other two protocols were intermediate in their performance in this regard (∼20% adapter dimers for NEXTflex and ∼10% for RealSeq). For matrices other than CSF, library quality was good and mostly adapter dimer-free for all protocols.

Crucially, the MDX protocol also demonstrated superior biomarker detection capabilities. It identified the highest number of unique miRNAs across every tested matrix (+69 for CCSN, +146 for FFPE, +119 for CSF versus the second-best benchmarked protocol; Figure 7B). Filtering criteria were applied to ensure the reliability of these findings: only miRNAs with RPM > 1 and mismatch-free read counts >= 5 were considered. This filtering, which on average eliminated around half of all identified miRNA species, aimed to remove likely artefactual reads, thereby enhancing the comparability of data. To confirm these findings operate independently of variations in effective sequencing depth (Figure 7C), we performed an *in silico* downsampling analysis. Randomly resampling reads to uniform depths across all libraries can isolate a protocol’s inherent detection sensitivity from raw sequencing output. Across all biological matrices, the MDX protocol consistently recovered the highest number of unique miRNAs at equivalent depth after a certain level of minimum depth was reached. This analysis confirms that the MDX protocol provides superior analytical sensitivity for comprehensive miRNome characterization in challenging sample types.

## Discussion

The precise analysis of the small RNA transcriptome especially in ultra-low-input biological matrices, such as CSF or exosomal fractions, is fundamentally limited by severe ligation biases and low sensitivity of standard library preparation methods ^21–23,36,37^. Standard commercial kits have a high background noise floor, may fail to produce libraries with enough reads mapping to species of interest, and frequently fail to capture lowly abundant and hard-to-ligate species in these difficult matrices. To resolve these bottlenecks, we developed the MDX protocol, which physically separates the 3’ and 5’ ligation steps for adapter-dimer minimization even with very low inputs and utilizes rationally engineered, structure-forcing 5’ adapters. The inherent challenge of precisely excising optically invisible intermediate products during PAGE purification, usually accomplished by using radioactivity, was resolved through the development of FLRs. These engineered, functionally inert markers enable precise, visual band excision without the hazardous or imprecise reliance on radiolabeling or simple estimation, significantly improving the reproducibility of the gel-excision step. Our final optimized protocol significantly outperforms leading commercial alternatives, reducing ligation bias on a highly adversarial synthetic miRNA panel (Figure 6A-B) and identifying more unique miRNAs in challenging matrices like CSF, FFPE, and CCSN (Figure 7B) while being able to prepare libraries from as little as 0.1 ng of total RNA equivalent of miRNAs (Figure S5C) despite relying on an intermediate gel purification step.

Rather than relying only on empirical screening to improve ligation performance, our approach was driven by bioinformatic analysis of the human miRNome. By profiling the MFE distribution of natural miRNA-adapter interactions, we engineered 5’ adapters designed to exceed a specific thermodynamic threshold to drive efficient cofold complex formation and by extension ligation. This targeted interaction overrides localized, unpredictable structural variations during the intermediate ligation step, reducing folding-related bias by standardizing the secondary structure of the 5’ ligation cofold complex. Specifically, our thermodynamic approach relies on calculating free energy at equilibrium. By modifying the 5’ adapter with engineered complementarity domains, we introduced predictable cofolding interactions. These interactions are designed to be strong enough to ensure near-complete target complex formation, even under conservative assumptions. In our optimal design identified, exemplified by the 5A3C6 adapter, the 5’ end engineered domain of the 5’ adapter binds to a site near the 3’ end of the 3’ adapter resulting in a stem-loop structure resembling a pseudo-circularized state in what we call “distal tethering”. Interestingly, the specific arrangement of structural domains proposed to be optimal in our study has not been studied in any of the prior structural works; instead, miRNA-specific or degenerate sequences were added to the 3’ end ^13,14^ or in the middle ^9^ of the 5’ adapter.

Our engineering concept is directly validated by our structural correlation analysis (Supplementary Figure S3), which reveals a clear shift in behavior between standard (Illumina TruSeq) and custom adapter architectures. The strong correlation between ΔMFE (our custom “driving force” metric) and bias for the native Illumina TruSeq adapters underscores how sequence-dependent thermodynamic variations drive severe representation bias. The collapse of these structural relationships in our custom engineered adapters (5A3C3 and 5A3C6) represents a likely case of "curse of success" – the global minimization of bias narrows the dynamic range to such a degree that any remaining variance becomes dominated by technical noise, such as the exactness of equimolar mixing in the synthetic panel. Interestingly, as the thermodynamic driving force increases further in the 5A3C3 variant, the correlation reverses even reaching statistical significance. This reversal may point to a state where excessively stable final biligated complex is introducing secondary structural barriers that inhibit subsequent enzymatic steps (in essence introducing a new type of bias), such as reverse transcription. However, given the small slope of this negative trend, its practical impact on overall library representation is likely minor in magnitude. Another interesting observation was the highly significant positive correlation between GC content and bias observed strictly for the baseline 5A3 adapter (ρ = 0.51). One possible explanation might be that high GC content might enable native miRNAs to partially "mimic" the structure-forcing tethering effect (due to higher potential for interaction strength) of our engineered domains.

Correlation analysis between cofold types and empirically measured library bias showed results that stand in contrast to prior reports ^9,15^, which attributed a large part of ligation bias to shifting distributions of favored cofold types through structural modifications of adapters. However, in our hands, this model had no predictive power in the case of modern Illumina TruSeq-based adapter sequences. Surprisingly, the highest total proportion of cofold types previously described as being favorable for T4 RNA ligase 1 (5, 9, and 13) ^9^ was observed for TruSeq adapters (∼82%) followed by 5A3C3 (∼70%), 5A3 (∼49%), and finally 5A3C6 (∼46%) (Supplementary Figure S3C). Despite these differences in distribution, for no adapter group did cofold types predict empirical bias (Supplementary Figure S3D). This discrepancy is likely caused by the differences in sequencing platforms utilized in their original work – Ion Torrent and an older version (V1.5) of Illumina small RNA adapters ^9,12,15^. Additionally, while Fuchs et al. ^9^ concluded that the sequence flanking the ligation junction is not the primary determinant of efficiency since degenerate sequences improved outcomes regardless of position, our findings point to a different model. We propose that modifications at different locations, even if utilizing the same method (i.e., degenerate bases), tackle bias through distinct mechanisms and that neither cofold type optimization nor junctional flexibility is sufficient to drive ligation bias minimization. Indeed, our baseline (5A3) and engineered designs (5A3Cx) incorporate degenerate junctional adapter nucleotides that are too short by themselves (2 nt) to appreciably increase adapter complementarity, yet such design can be significantly improved with engineered complementarity domains at the opposite end of the adapter sequence. This, in conjunction with the cofold type analysis discussed earlier, also implies that our method of flattening the thermodynamic landscape directly without requiring a major favorable redistribution of cofold types or introducing more sequence degeneracy provides a unique source of bias minimization. Based on the above, we suggest that optimization strategies must be layered and executed in an adapter-sequence and platform-specific manner to minimize the overall bias of small RNA library preparation workflows.

It should be noted that our engineered adapters exhibited ligation efficiencies much higher than the original unmodified baseline, a unique benefit with respect to protocol sensitivity, and this effect was not directly correlated with ligation bias reduction (Figure 3). In connection with this observation, we theorize that our engineered complementarity domains increase effective local concentration of reactants. In ultra-low-input biological matrices, such as CSF, dilute analyte concentrations severely limit the kinetic frequency of productive intermolecular collisions. It is likely that distal tethering artificially increases the effective local concentration of the reactive 5’ adapter at the ligation site, effectively converting a kinetically limited intermolecular reaction into a more efficient intramolecular-like process. The existence of such a mechanism is consistent with a couple of observations. First, ligation yield and bias (Figure 3A-B) show different underperformers – 5A3C4 for yield versus 5A3C2 for bias – highlighting a clear decorrelation between reaction efficiency and ligase substrate preferences. This is also supported by the observation that junctional degenerate bases by themselves (5A3) were insufficient to drive high 5’ ligation conversion yields (Figure 3A), even though the baseline ligation bias for this adapter was comparably restrained in absolute terms (Figure 3B). And second, the only reaction parameter we internally found to improve 5’ ligation was increasing its duration, an effect consistent with a collision frequency limited reaction (Figure 4C). While we could not definitely confirm that this mechanism is the primary one, it is clear it does not only act to enable substrate-specific ligase activity, but it also drives ligation closer to completion even for species with low baseline bias.

Apart from improving ligation performance, our proposed adapter design has an additional advantage. The structural simplicity of our optimized 5A3C6 adapter architecture natively supports the integration of UMIs (Supplementary Figure S2) unlike approaches where longer degenerate sequences of nucleotides are the primary means of ligation bias reduction (as they inherently reduce the bit entropy of the sequence being they are supposed to interact with other oligonucleotides in a sequence-specific manner). While not strictly required for standard relative quantification and differential gene expression workflows, this built-in compatibility provides a critical tool for ultra-low-input discovery scenarios where correcting for reverse transcription and PCR amplification duplication artifacts might be necessary to maximize accuracy ^31^.

Intriguingly, this UMI sequence is in principle very similar to the approach used in ^9^ to improve ligation bias and yields; yet, we saw no evidence of it affecting final ligation bias when combined with our engineered domains (Supplementary Figure S2B-C) which is consistent with these degenerate bases showing an expected, mostly random distribution of nucleotides per position (Supplementary Figure S2D-E). This might suggest that either our approach and theirs share a mechanism of action that is already maximized by presence of one of them or that our engineered domains override it.

Despite the advantages of our proposed MDX workflow, we acknowledge certain limitations in the present study. First, the MDX protocol is inherently more laborious and lower-throughput than standard commercial workflows. The requirement for intermediate PAGE purification, while essential for physically separating the 3’ and 5’ ligation steps to minimize adapter-dimer formation, limits its applicability for high-volume routine sequencing. However, this trade-off is a natural consequence of designing a protocol specifically for small RNA biomarker discovery, where ultimate sensitivity and representational correctness are prioritized over throughput, and where downstream validation is typically performed using targeted, higher-throughput RT-PCR assays.

Second, our evaluation of the engineered 5’ complementarity domains may understate their independent effect. Because our baseline protocol (using the 5A3 adapter) already incorporated degenerate junctional bases – a feature known to significantly reduce bias ^13,14^ – the relative bias reduction achieved by the final 5A3C6 variant appears modest (∼20% relative decrease), despite achieving an excellent absolute coefficient of variation. The simultaneous improvement in final product yield (∼80% increase), however, highlights the multifaceted benefits of this synergistic approach.

While highly optimized, bias-reducing workflows like the MDX protocol may not be strictly necessary for routine differential expression analysis of moderately-to-highly abundant small RNAs with good ligation efficiencies. However, mitigating quantification bias is a crucial requirement for biomarker discovery. In these scenarios, knowing the true biological abundance is critical, as artificially underrepresented molecular species might otherwise be overlooked as viable clinical targets. Additionally, highly biased protocols are susceptible to unpredictable representational shifts caused by compositional changes in the input, such as varying ratios of background nucleic acids. By minimizing representational bias and maintaining high sensitivity in challenging matrices, the MDX protocol offers a robust solution for high-confidence miRNome characterization when this extra robustness is required.

## Materials and Methods

### Materials

All standard oligonucleotide synthesis reagents (Deblock, Cap A, Cap B, Activator, Oxidizer, Acetonitrile for DNA synthesis) were purchased either from emp Biotech or Merck. Other reagents used: DMT-protected DNA phosphoramidites with base-labile protecting groups (Glen Research, US; Hongene, CN), DMT; TBS-protected RNA phosphoramidites with base-labile protecting groups (Glen Research, US; Hongene, CN), PT-Amino-Modifier C6 CPG solid glass support (Glen Research, US), Amino-Modifier C6 dT phosphoramidite (Glen Research, US), Solid Chemical Phosphorylation Reagent II (Glen Research, US), AF488 NHS ester (Lumiprobe, DE).

### Oligonucleotide synthesis and purification

All oligonucleotides used in this project were synthesized on a Dr. Oligo 48 (Biolytic, US) synthesizer on a 1.0 μmol scale. Synthesized oligonucleotides were deprotected according to the instructions of the manufacturer of phosphoramidites (Glen Research, US). Sequences of all oligonucleotides used in this study can be found in Supplementary Table 1 and Supplementary Table 2.

For regular DNA or RNA oligonucleotides, crude oligonucleotides were purified either using denaturing polyacrylamide gel electrophoresis (DMT-OFF synthesis) or using Glen-Pak cartridge purification (Glen Research, US; DMT-ON synthesis). The resulting purified oligonucleotides were desalted, dried and subjected to quality control using denaturing polyacrylamide gel electrophoresis as previously described ^38^.

Phosphorylated RNAs for the synthetic miRNA panel (described below) were purified using a Shimadzu HPLC system equipped with a CMB-40 system controller, LC-40D solvent delivery system and a SPD-M40 UV-Vis detector. The purification was done in an ion-exchange mode using a DNAPac PA-100 column (ThermoFisher, US); 1 ml/min, gradient of 1.5 M NaCl in 100 mM TEAA (pH = 7.0); 0 -> 100% in 35 min. The resulting purified RNA was further desalted using the same HPLC setup utilizing a BabyBio DSalt 5 ml column (Bio-Works, FI), eluting with HPLC grade water (Sigma-Aldrich, US). The final product was analyzed using denaturing polyacrylamide gel electrophoresis identically to regular oligonucleotides as described above.

In case of amino-modified oligonucleotides, the pure precursors were post-synthetically labelled using AF488 NHS ester following a previously described procedure ^38^. For the preadenylated 3’ adapter, the pure 5’-phosphorylated adapter precursor was adenylated and further purified as described previously ^39^. The final fluorescently labeled and preadenylated sequencing adapter was analyzed using mass spectrometry (ESI-MS) to confirm its identity and purity.

### Samples

Technical evaluation of the miRNA sequencing protocols was performed using three human matrix types: commercial FFPE breast cancer tissue, CSF, and cerebral organoid CCSN. FFPE samples were obtained from AMSBio (US). Pooled, de-identified CSF and cerebral organoid CCSN samples were provided by the ADDIT-CE consortium via collaborators at St. Anne’s University Hospital (FNUSA) and Masaryk University in Brno, respectively. Informed consent and ethical approvals pertaining to CSF sample use were obtained and managed directly by FNUSA as part of the long-running Czech Brain Aging Study in accordance with their institutional review board guidelines and the Declaration of Helsinki. Cerebral organoids of cultures generated using previously established, fully de-identified human cell lines were grown and their supernatants collected as described in ^40^. CSF and CCSN were stored at -80 °C while FFPE specimens were stored at 4 °C until sectioning and library preparation. All biomedical research procedures were performed in compliance with the approval issued by the Regional Public Health Authority of Bratislava (RÚVZ Bratislava, Slovakia), approval reference RÚVZBA/OPPL/7943/14225/2023, and in accordance with applicable national regulations.

### RNA input preparation

Total RNA was isolated from both tissue and biofluid samples to evaluate MDX protocol performance across diverse input types. FFPE samples were extracted using the NucleoSpin totalRNA FFPE XS kit (Macherey–Nagel, DE). CSF and CCSN samples were extracted using the Urine microRNA Purification Kit (Norgen Biotek, CA). For comparing protocol performance on biological matrices, inputs were standardized either by RNA mass (100 ng FFPE RNA) or by input volume for biofluids (400 µl CSF and 1000 µl CCSN). For each sample type, 10 individual samples were pooled prior to RNA extraction to generate a single composite input. This pool was used uniformly across all library preparation workflows to ensure identical starting material for cross-protocol comparison. For FFPE samples, RNA quantification was performed using the Qubit RNA High Sensitivity assay (Invitrogen, US). For protocol benchmarking experiments, libraries were generated using either equimolar synthetic miRNA pools or biological total RNA samples. Synthetic inputs consisted of either an 88x miRNA pool or an expanded 150x miRNA equimolar panel at either 0.7 pmol per reaction (experiments involving gel-based visualization) or to an amount corresponding to target “total RNA equivalent” input. This term defines a quantity of pure synthetic miRNA equal to the amount of endogenous miRNA fraction naturally contained within a given amount of total RNA from a true biological specimen. This relationship was determined empirically by assaying libraries made from high-quality, adapter-dimer-free, high-input biological libraries (1000 ng of FFPE breast cancer total RNA with high PCR efficiency and yielding a single, expected product) by RT-PCR and comparing it against a calibration curve derived from similarly pure libraries constructed from a known concentration of our synthetic miRNA panel. The conversion factor derived from this difference was then used to dilute the synthetic miRNA panel stock to approximate true miRNA content in biological specimens of a target input amount (0.1 – 1000 ng; 0.55 – 5500 amol).

### Library construction

For the 3′ ligation step, reactions were assembled to final concentrations of 15% PEG 8000, 20% DMSO (omitted in the unoptimized baseline workflow), 1x T4 RNA Ligase Reaction Buffer, and 100 nM (1 pmol) 3′ adapter (3AB1), together with the RNA input sample. Reactions were denatured at 90°C for 1 min and cooled on ice for 2 min. T4 RNA Ligase 2 truncated KQ (100 U final; NEB M0373) and RNaseOUT™ (20 U final; Invitrogen) were then added to a final reaction volume of 10 µl, followed by incubation for 4 h at 25°C. Following 3′ ligation, reactions were supplemented with fluorescent ligation rulers (FLR18 and FLR25; 0.4 pmol each), denatured at 95°C for 1 min, and cooled on ice for 2 min. An exception was made when evaluating ligation optimization experiments and evaluating FLR ligation bias interference where FLRs were omitted in certain cases. To minimize 3′ adapter carryover, 3′ ligation products were separated by 15% denaturing urea-PAGE for 3 h at a constant 400 V, with maximum power limited to 15 W. FLR markers co-migrated with corresponding ligated miRNA species, enabling precise excision of target products between the 18 nt and 25 nt fluorescent boundaries. Gels were visualized and excised using the iBright™ FL1500 Imaging System (Invitrogen, US) in band excision mode. Excised gel fragments were pulverized with a disposable pestle and submerged in 0.3 M NaCl. Gel suspensions were subjected to three freeze–thaw cycles (−80°C for 15 min and 35°C for 3 min), followed by overnight elution with rotation at 15 rpm at 4°C. Eluted material was separated from gel debris using Spin-X® centrifuge tube filters containing 0.45 µm cellulose acetate membranes (Corning, US), followed by ethanol precipitation using GlycoBlue™ coprecipitant (Invitrogen, US).

Recovered 3′-ligated RNA products were resuspended in 2.5 µl nuclease-free water and transferred to a 5′ ligation reaction containing 15% PEG 8000, 1x T4 RNA Ligase Reaction Buffer, 1 mM ATP, and 50 nM (0.5 pmol) 5′ adapter of total volume 10 µl. The final optimized workflow used adapter variant 5A3C6, whereas optimization experiments evaluated additional variants including 5A3, 5A3C1–5A3C7, and 5A3C6U. Reactions were denatured at 90°C for 1 min and cooled on ice for 2 min prior to addition of T4 RNA Ligase 1 (10 U final; NEB M0204) and RNaseOUT™ (20 U final). Final reactions were incubated for 4 h at 37°C (1 h in the unoptimized baseline workflow). Biligated products were then purified using the Oligo Clean & Concentrator kit (Zymo Research, UK).

Commercial small RNA library preparation workflows used during protocol benchmarking – TruSeq Small RNA Library Prep Kit (Illumina, US), NEXTflex Small RNA Sequencing Kit v4 (Revvity, US), and RealSeq®-AC Small RNA Library Preparation Kit (RealSeq Biosciences, US) – were performed precisely according to manufacturer instructions up until the cDNA step. For the MDX protocol, purified biligated products were pre-heated with RTP primer (50 pmol) and dNTPs (0.5 mM final concentration) for 5 min at 65°C and then cooled on ice for 2 min. First-strand cDNA synthesis reactions (20 µl total volume) were completed by adding 1x SSIV buffer, SuperScript™ IV reverse transcriptase (200 U final; Invitrogen, US), RNaseOUT™ (40 U final), and 5 mM DTT, followed by incubation at 56°C for 30 min and 80°C for 10 min. From this stage onward, cDNA products from all workflows underwent identical downstream amplification and purification procedures. PCR amplification reactions (25 µl) contained 1x KAPA HiFi HotStart ReadyMix (Roche, CH), 1 µM SYTO 9 fluorescent nucleic acid stain (Invitrogen, US), 10-fold diluted cDNA template, 5AP1 forward primer (12.5 pmol), and RPI-ex10 (1–48) indexed reverse primers (12.5 pmol). Two identical PCR reactions were prepared for each sample. The first aliquot (1/2) was amplified to plateau phase to determine Cq values, whereas the second aliquot (residual cDNA) was amplified to exactly Cq + 5 cycles using an AriaMx Real-Time PCR System (Agilent, US). Cycling conditions consisted of initial denaturation at 95°C for 5 min followed by up to 35 cycles of 95°C for 5 s and 62°C for 10 s for denaturation and annealing/extension, respectively.

Amplified libraries were resolved by 15% native PAGE at 200 V for 1 h followed by 400 V for 2 h, with maximum power limited to 15 W. Gels were post-stained using 1× SYBR Gold (Invitrogen, US), and target library bands were excised and recovered using the same gel extraction and precipitation procedure used for products after 3’ ligation but omitting the 3 freeze thaw cycles described previously. Following precipitation, final library pellets were resuspended in elution buffer (10 mM Tris-HCl, pH 8.5, 0.1% Tween-20). Library quality control was performed using the 4150 TapeStation system (Agilent, US) with High Sensitivity D1000 ScreenTape assays to verify fragment size distribution and determine molarity prior to library pooling and sequencing on an Illumina NextSeq 2000 platform. Libraries were pooled equimolarly to a final loading concentration of 800 pM and sequenced in a 100 bp paired-end configuration on the NextSeq 2000 platform (Illumina, US). Sequencing was performed with a target output of 10 million reads per sample.

### Bioinformatic analysis

#### Sequencing data trimming and mapping

Bcl2fastq (v2.20.0) was first used to demultiplex and convert BCL basecall files from the sequencer to fastq files. Next, sequencing library reads were trimmed and mapped against a reference database using MiRACLE, an in-house tool designed for advanced trimming and seed-less, fuzzy, global sequence alignment of short reads against small-to-medium sized annotation databases like miRbase (v22.1) ^34^. This tool also generates a feature count table for each processed sample and extracts custom adapter parts (such as UMIs, junctional degenerate bases, etc.) grouping them by core mapped sequence reads for downstream analyses. Results generated by MiRACLE are broadly comparable with using Cutadapt ^41^ and Bowtie ^42^ with Pearson correlation of feature count tables exceeding 0.995 for synthetic miRNA panel inputs and 0.9 for biological specimens (data not shown).

The adapter sequences were trimmed if their sequence was with a maximum of four errors permitted. If degenerate bases were incorporated at the 3’- or 5’-ends of the sequenced fragments, these were also removed. Filtering step removed low-quality reads (mean Phred quality score < 20 or Phred quality score of any base < 5 after trimming) or reads that were longer than 28 bp or shorter than 16 bp except for reads shorter than 8 bp that were instead categorized as adapter dimers. Maximum of two errors were permitted for mapping to reference. For different experiments the reference used was either list of known miRNAs in the synthetic panel mix used for a given experiment or miRbase.

For samples consisting of synthetic equimolar miRNA panels, no further processing was performed, and feature count tables from the first phase were used for analyses directly. However, for biological samples (e.g. CCSN, FFPE, CSF), primary analysis by MiRACLE was followed by alignment to the human genome using Bowtie (v.1.0.0).

Trimmed reads mapping perfectly to miRBase sequences were excluded from this further processing, as they were considered high-confidence miRNA matches. All remaining reads not categorized as adapter dimers or filtered out were collapsed and aligned to the primary assembly of human genome GRCh38 (v.113) using Bowtie. Only alignments with at most 2 mismatches (in the first 18 bp seed region of the read) were considered. Single best alignment is reported for reads with no more than 30 possible alignments. The genomic index used for Bowtie alignment was supplemented with an additional sequence of a complete rRNA 45S cluster obtained from RefSeq (accession KY962518.1), as annotation of genes on this cluster is absent from ENSEMBL due to their localization within repeat regions, making them difficult to assemble. The feature annotation file therefore incorporates all ENSEMBL features (v.113), the three rRNA 45S components (5.8S, 18S and 28S) and high confidence set of 432 tRNAs from GtRNAdb v.2.0.

Feature count tables were generated from reads aligned to either a miRbase sequence using MiRACLE or an exonic region of a genomic feature with Bowtie. In instances where read was mapped to both, the alignment with the lower edit distance was retained.

Reads mapped to different features with identical edit distances were considered ambiguous and discarded. To prevent redundancy, miRNA features identified by Bowtie were renamed from ENSEMBL IDs to their corresponding miRBase name, if possible.

The final read count table comprises perfect miRBase hits identified by MiRACLE and resolved, unambiguous hits identified by the Bowtie-MiRACLE comparison.

The final read count tables include raw counts, perfect raw counts, RPM. Perfect raw counts refer to the number of reads mapping with zero errors. RPM is calculated as fraction of raw counts from each million of mapped miRNA reads within the library. To remove likely artifactual or highly noisy hits, miRNA feature counts with an RPM < 1 and perfect raw counts < 5 were excluded from subsequent analyses including to define presence/absence for visualization purposes.

### Synthetic miRNA panel design

To systematically evaluate ligation bias and protocol performance, we developed custom synthetic human miRNA equimolar panels. An initial 88x panel was generated for early pilot optimizations and analytical characterizations, followed by a comprehensive 150x panel used for most experiments. Panel composition was determined using a custom Python script designed to sample sequences from miRBase (v22.1). The selection algorithm was seeded with well-known and highly expressed miRNAs to ensure biological relevance, and the script used simulated annealing to iteratively adjust a subset of miRBase miRNAs that accurately mirrors the structural complexity of the human miRNome. Specifically, the objective was to minimize the Kullback-Leibler divergence between miRBase and target distributions of sequence length, GC content, predicted heterodimer interactions with other miRNAs in the pool (determined using BioPython’s Aligner using default settings in ‘local’ mode), and self-folding strength (computed using ViennaRNA as detailed in “Thermodynamic and structural modeling of adapter-miRNA interactions”; all weights of 1.0) while concomitantly maximizing total sequence entropy (weight of 0.5). Furthermore, among initial seeds were also included sequences with hand-selected "extreme" structural characteristics, miRNAs reported to have poor ligation efficiencies in previous studies or white papers, as well as pairs predicted to form exceptionally strong heterodimers. The final composition of synthetic miRNA panels can be found in Supplementary Table 1.

### Downstream data analysis

All data analysis was performed using custom Python (3.12.11) scripts based on using standard scientific libraries, *scipy* (1.16.2) and *numpy* (2.1.2).

Ligation bias for libraries generated from equimolar synthetic miRNA panels was quantified by comparing the observed read count for each miRNA to its expected theoretical read count. Because the synthetic panels consisted of equimolar mixtures, the expected read count for any individual miRNA was determined by scaling its theoretical count (identical for all miRNAs) to the effective sequencing depth. Bias was subsequently expressed as the log2-transformed ratio of observed to expected read counts.

To evaluate the similarity of ligation bias profiles across engineered 5’ adapters, hierarchical clustering was performed using the ‘average’ linkage method and the ‘euclidean’ distance metric.

For cross-protocol performance comparisons in biological matrices, the overlaps of detected miRNA subsets between the evaluated protocols were visualized as four-way Venn diagrams using the *pyvenn* library.

To isolate the inherent analytical sensitivity of the protocols from variations in total sequencing depth, an *in silico* downsampling analysis was performed. The numpy.random.multinomial() function was applied to feature count tables to simulate uniform read depths.

To analyze the randomness of UMI region in the 5A3C6U adapter and quantify sequence-specific ligation bias, we calculated the position-specific Shannon bit entropy for the UMI nucleotides. For each position within the UMI, the observed entropy was calculated across all reads mapped to a specific miRNA. Because a perfectly random, equiprobable distribution of four nucleotides yields a maximum theoretical entropy of 2 bits, sequence-specific bias was expressed as the deviation from this value representing absolute randomness (value close to 0 indicates a highly random distribution, whereas higher values indicate sequence bias). For the sequence logo ^43^ graph generation, *logomaker* library ^44^ was used on averaged data across all miRNA species.

To ensure reproducibility, a fixed random seed of 42 was applied across all scripts utilized in this study.

### Sequencing library composition and biotype aggregation

To evaluate the distribution of sequence reads, mapping annotations were collected and grouped into defined biotype categories prior to relative abundance analysis.

Sequences assigned to protein-coding sequences, unannotated intergenic features (no feature), transcripts of unknown environmental origin (TEC), ribozymes, sequencing artifacts, and all pseudogenes (including processed, unprocessed, unitary, and immunoglobulin or T-cell receptor pseudogenes) were combined into a merged Genomic category. Transfer RNA (tRNA) and ribosomal RNA (rRNA) classes included both nuclear and mitochondrial variants (Mt_tRNA and Mt_rRNA), as well as ribosomal pseudogenes. Small non-coding transcripts including small nuclear RNA (snRNA), small nucleolar RNA (snoRNA), small Cajal body-specific RNA (scaRNA), small cytoplasmic RNA (scRNA), vault RNA, small RNA (sRNA), miscellaneous RNA (misc_RNA), and long non-coding RNA (lncRNA) were merged into a single category designated as Other ncRNA. The final percentage of reads attributable to miRNA, tRNA, rRNA, Other ncRNA, Genomic, special sequences (external calibrators ^27^ – used only for FFPE samples, FLR fragments, and Illumina StopOligo sequence), adapter dimers, and unmapped reads was calculated by normalizing the count of each category against the total read depth of the respective library.

### Thermodynamic and structural modeling of adapter-miRNA interactions

To compute an approximation of the driving force for the structural conversion of the 5’ adapter strand into the target heterodimeric 5A & miRNA-3A complex, we calculated the MFE difference as:

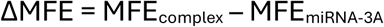

where MFE_miRNA-3A_ is the self-folding minimum free energy of the first ligation product, and MFE_complex_ is the cofolding minimum free energy of 5’ adapter and the first ligation product.

We established a conservative thermodynamic threshold (MFE_gap_) to guide the design of the complementarity domains added to the 5’ adapter. This threshold represents the energy gap required to achieve a targeted complex formation rate at equilibrium, maintaining ΔMFE ≤ MFE_gap_ for most miRNAs considered. The threshold is based on the standard free energy change ΔG^◦^:

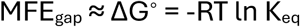

where R is the gas constant, T is temperature in Kelvin, and K_eq_ is the equilibrium constant. The equilibrium constant is defined by the concentrations of the interacting species:

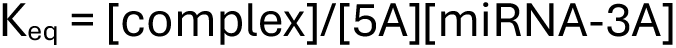

where [complex] is the cofolding heterodimer concentration, [miRNA-3A] is the concentration of the first ligation product, and [5A] is the 5’ adapter concentration in the ligation reaction. Because the adapter strands are present at a massive excess relative to the target miRNA species, the free adapter concentration [5A] was treated as a constant under a pseudo-first-order approximation. Setting a target conversion threshold of 95% simplifies the equilibrium constant to K_eq_ = 19 / [5A]. Utilizing a 5’ adapter concentration of 50 nM and RT = 0.616 kcal/mol (at 37°C), the final MFE_gap_ of -12.2 kcal/mol established our conservative engineering threshold used to design complementarity domains to be added to the 5’ adapter.

To compute MFE values and secondary structures, ViennaRNA’s ^30^ RNAcofold function was used. Model parameters used were as follows: Turner 2004 RNA thermodynamic parameters ^45^, 37°C, 0.04 M Na^+^ (derived from a 0.05 M Tris buffer at pH 7.5, reflecting 80% ionization), 0.01 M Mg^2+^ (converted to Na^+^ equivalents using the equation 3.3 x √[Mg^2+^] and summed with [Na^+^]), and with “noLP” flag set to true. With RNA thermodynamic parameters, the predicted interactions involving the DNA-only 3’ adapter would be inherently biased towards more negative MFEs and more complex secondary structures, as ViennaRNA does not support chimeric RNA:RNA/DNA interaction modeling. To partially alleviate this issue, we used the soft constraints feature of ViennaRNA where specific nucleotide index pairs had an energy penalty of +0.5 kcal/mol applied, the average difference between RNA:DNA ^46^ and RNA:RNA ^45^ base pairing nearest-neighbor parameters, to penalize chimeric interactions as a rough proxy for the weaker nature of RNA:DNA bonds. Additionally, to maintain a conservative estimate of the driving force for conversion and to avoid modeling complex competitive interactions (which act as thermodynamic sinks in complex biological mixtures) with other nucleic acids present during ligation, we deliberately omitted 5’ adapter self-folding and restricted cofolding analysis to inter-molecular interactions only. For structural designs containing degenerate bases, all degenerate positions (Ns) were expanded to a complete set of possible sequence variants, and the variant yielding the worst MFE was selected to model the worst-case scenario. The only exception was when investigating structural relationships between empirically measured bias and structural features when the empirically most common degenerate base combination was used instead.

For structural studies investigating relationships between cofold types and bias, every pair of sequences was classified into 1 of 16 cofold types as originally proposed in ^15^.

This was performed using a custom function written in Python that used the dot-bracket notation output of the ViennaRNA cofold function as its input.

### Densitometry

Ligation efficiencies and elution recovery were quantified via densitometric analysis of polyacrylamide gels. Visual detection of oligonucleotide bands was enabled by fluorescent labels covalently attached to the 3′ adapter, whereas input RNA and the 5′ adapter remained unlabeled. Following gel electrophoresis, fluorescent signal intensities were captured using an iBright™ FL1500 Imaging System (Invitrogen, US) and analyzed using the Fiji ^47^ distribution of ImageJ software (version 2.16.0 / 1.54p). For each lane, individual band intensities were quantified by measuring the integrated density of pixels within identically sized regions of interest, with local background subtraction applied uniformly across all analyzed lanes.

Ligation yield was expressed as a percentage of the ligation product band intensity relative to the cumulative signal intensity of all bands within the same lane. If the fluorescently labeled component of the ligation reaction was present in excess relative to the unlabeled component, the calculation was adjusted to account for this fact by subtracting a percentage from the total equal to the excess (detailed in the respective figure descriptions). Similarly, elution yield was expressed as a percentage of the gel-purified ligation product band intensity relative to the unpurified fraction intensity.

### Statistics and data presentation

Statistical analyses were performed using custom scripts written using the libraries *numpy* and *scipy* in Python. Correlations between structural features and empirical bias were evaluated using the Spearman rank-order correlation coefficient. To assess technical reproducibility, inter-replicate consistency, and inter-protocol comparisons, Pearson correlation coefficients were calculated using log10-transformed normalized read counts (RPM + 1). Comparisons of empirical bias vs cofold type distribution across multiple adapter designs or protocols were assessed using the Kruskal-Wallis test. In all box plots presented, the center line indicates the median, and the bounding box (if present) represents the interquartile range. Specific technical replicate counts are detailed in the respective figure legends.

## Acknowledgements

The authors gratefully acknowledge our ADDIT-CE consortium partners, including the Czech Brain Aging Study at St. Anne’s University Hospital (FNUSA, Brno, Czechia) for providing CSF samples, and Dr. Dáša Bohačiaková (Masaryk University, Brno, Czechia) for providing the cerebral organoid CCSN samples. We also thank Geneton s.r.o. (Bratislava, Slovakia) for their assistance in performing high-throughput sequencing. This project has received funding from the European Union’s Horizon Europe research and innovation programme under grant agreement No 101087124, EU NextGenerationEU through the Recovery and Resilience Plan for Slovakia under the project No. 09I01-03-V04-00011, and the International Centre for Genetic Engineering and Biotechnology ICGEB No. CRP/SVK22-04_EC. Gemini (version “3.1 Pro”, Google, US; https://gemini.google.com/) was used to improve the clarity and grammar of the manuscript. A few graphical icons used in the Figure 1 schematic were also generated using Gemini. For figures displaying oligonucleotide secondary structures, initial layouts were generated utilizing the RNAstructure software package ^48^ and were subsequently modified in InkScape (1.4.3). All outputs were carefully reviewed, revised, and integrated by the authors, who take full responsibility for the final content.

## Author contributions

A.S. and P.Č. conceptualized and planned the study. N.D. and A.S. conducted bioinformatic analyses. A.S. designed engineered adapters and wrote the synthetic miRNA panel generation script. J.M., S.B., I.Č., and M.Č. synthesized and purified all oligonucleotides used in the study. S.A.B. and D.L. performed all wet-lab experiments including sequencing library preparation, workflow optimization, and gel experiments.

A.S., N.D., S.A.B., D.L., and P.Č. verified the underlying data, analyzed, and interpreted the data, prepared figures, and wrote the manuscript. All authors had full access to the data and provided feedback on the manuscript.

## Data availability

All raw and processed sequencing data generated during this study, as well as the underlying data for the figures presented, are available from the corresponding authors upon reasonable request. The full sequencing datasets will be deposited into a public repository prior to formal peer-reviewed journal submission.

## Code availability

The custom Python scripts utilized for generating the synthetic miRNA panels and for computing MFE values are available from the corresponding authors upon reasonable request. The proprietary joint trimming-mapping tool used in this study, MiRACLE, is currently undergoing internal benchmarking and further development and is not publicly available at this time. Future revisions of this manuscript will incorporate analyses conducted using standard, open-source bioinformatic pipelines (e.g., Cutadapt combined with Bowtie).

