## Supplementary Data for "Fluorescently guided workflow with rationally engineered 5′ ligation adapters for high-sensitivity and low-bias small RNA sequencing"

**S1A**

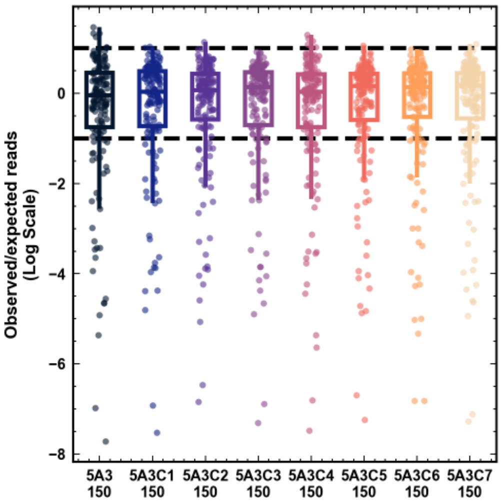
Observed/expected reads (Log Scale)
0
−2
−4
−6
−8
5A3 150
5A3C1 150
5A3C2 150
5A3C3 150
5A3C4 150
5A3C5 150
5A3C6 150
5A3C7 150

**S1B**

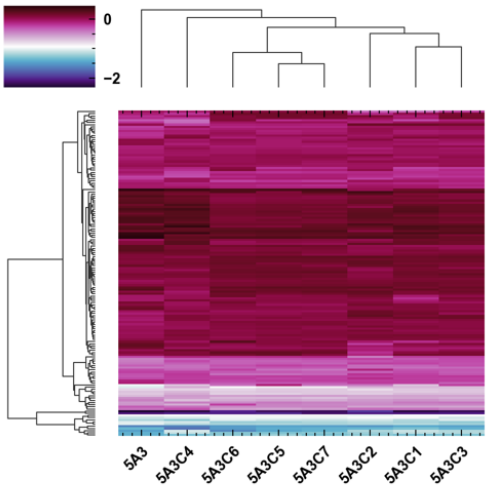
0
−2
5A3
5A3C4
5A3C6
5A3C5
5A3C7
5A3C2
5A3C1
5A3C3

**Supplementary Figure S1. Ligation bias profiles and clustering analysis of engineered 5' adapters.**

A) Boxplots depicting ligation bias (average of 2 technical replicates), expressed as the log2-transformed ratio of observed to expected read counts for the baseline 5A3 adapter control and modified custom variants 5A3C1 through 5A3C7 using the 150x synthetic miRNA panel. Numbers underneath group labels indicate number of unique miRNAs detected for individual groups. Horizontal dashed lines define the +/- 1 log2 fold change low-bias range.

B) Hierarchical clustering analysis of per-miRNA ligation profiles across tested designs, demonstrating that custom-engineered adapters incorporating a complementary domain at their 5' terminus cluster tightly together.

## S2A

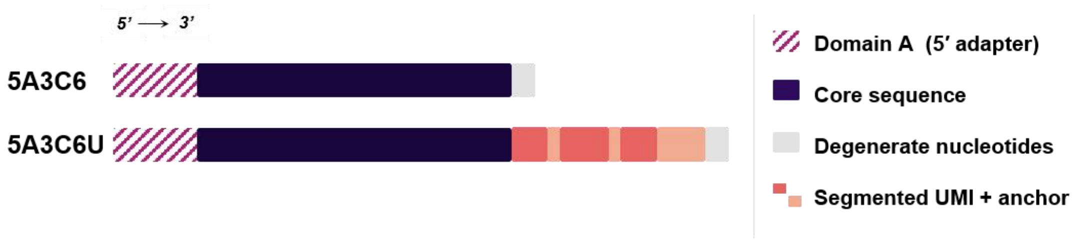


## S2B

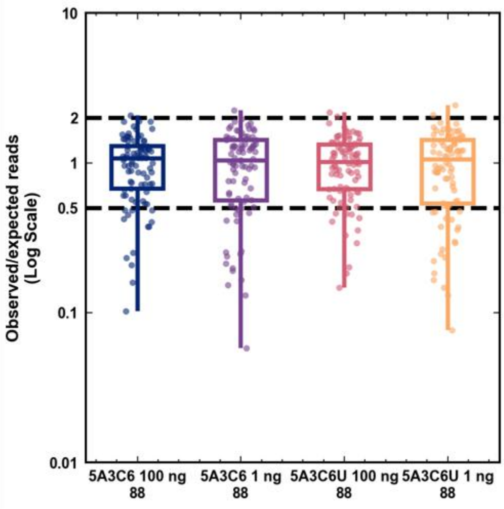


## S2C

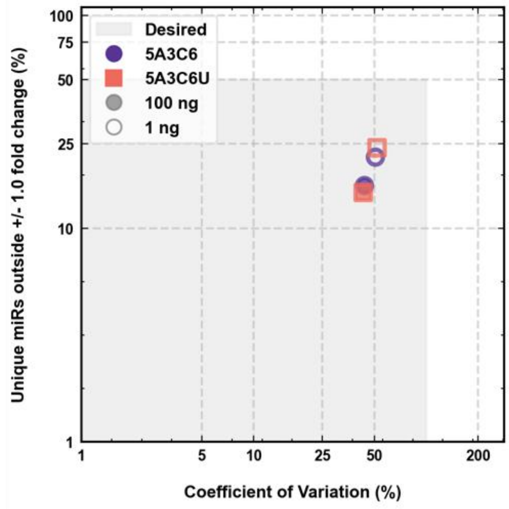


## S2D

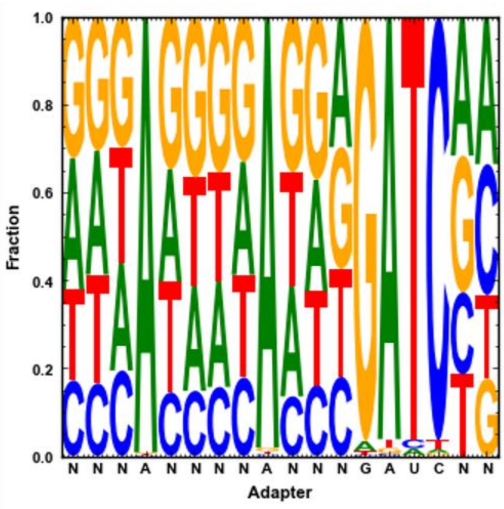


## S2E

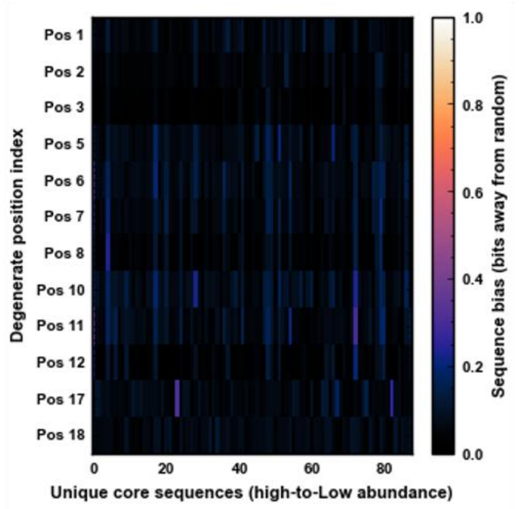

**Supplementary Figure S2: Structural design and validation of UMI-containing 5' adapters.**

A) Schematic comparison of the optimized 5A3C6 design and its UMI-modified derivative, 5A3C6U. The baseline 5A3C6 adapter consists of a 5'-terminal, 3'-adapter-binding domain (‘A’) followed by an Illumina-compatible core sequence and terminal degenerate bases. In the 5A3C6U variant, a segmented Unique Molecular Identifier (UMI) and its associated anchor sequence are integrated directly between the core sequence and the 3'-terminal degenerate nucleotides.

B) Boxplots depicting ligation bias (1 technical replicate), expressed as the log2-transformed ratio of observed to expected read counts, comparing 5A3C6 (no UMI) and 5A3C6U (with UMI) performance at 100 ng and 1 ng total RNA equivalent inputs using an 88x synthetic miRNA panel. Numbers underneath group labels indicate number of unique miRNAs detected for individual groups. Horizontal dashed lines define the +/- 1 log2 fold change low-bias range.

C) Scatter plot of above summarizing protocol performance by tracking the percentage of unique miRNAs falling outside a +/- 1 log2 fold change range against the overall Coefficient of Variation (CV, %), showing no obvious differences between adapters with or without UMI incorporation across all tested configurations.

D) Sequence logo mapping the relative fraction and composition of individual nucleotides across positions within the UMI and anchor regions of the engineered adapter, demonstrating broadly random distribution for UMI degenerate nucleotide positions (first 10 Ns from left-to-right). Junctional degenerate bases show a different profile consistent with their role in mitigating ligation-specific junctional preferences.

E) Heatmap evaluating sequence bias (expressed as bit entropy per position deviation from randomness) calculated across degenerate position indices for unique core sequences sorted from high to low abundance. For most sequences no bias can be seen suggesting that near full bit entropy of the UMI is available for deduplication and that these degenerate bases do not contribute and/or override the proposed ligation reduction mechanism of the 5’ engineered domain.

## S3A

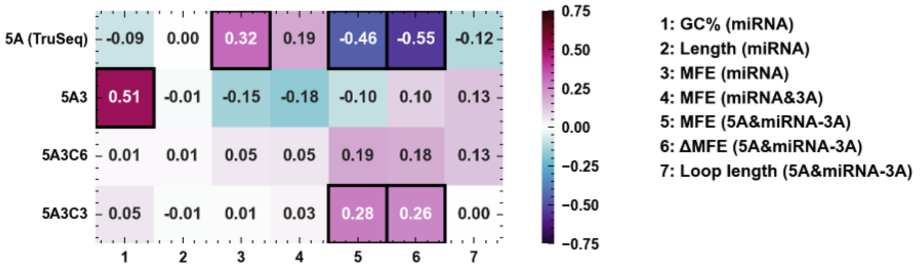


## S3B

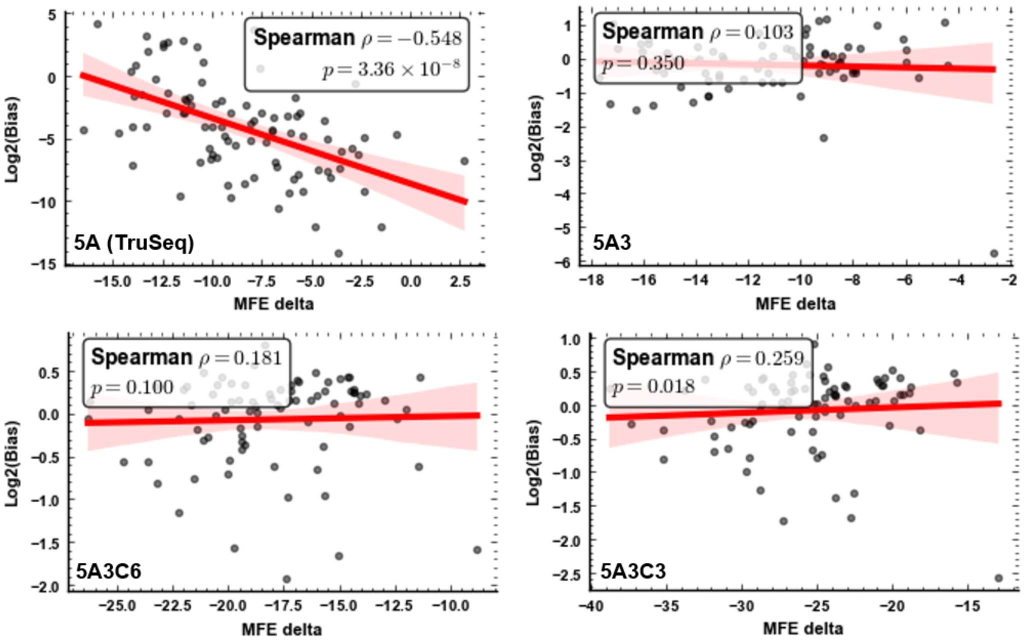


## S3C

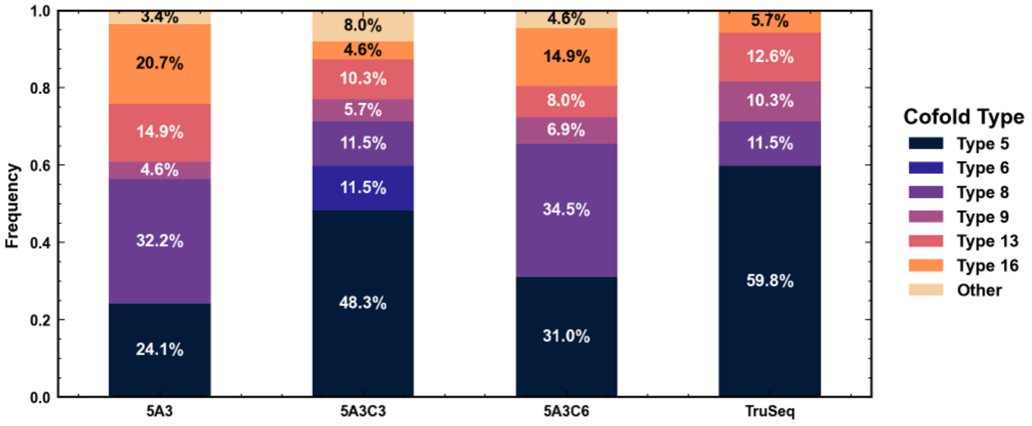


## S3D

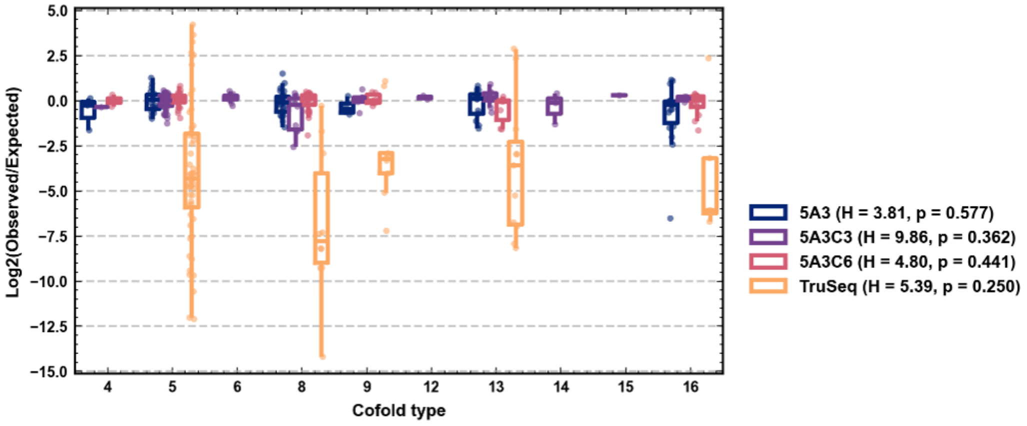

**Supplementary Figure S3. Thermodynamic and structural correlation analysis across baseline and engineered adapters.**

A) Heatmap summarizing Spearman correlation coefficients (ρ) calculated between specific miRNA structural features - GC content, sequence length, native miRNA MFE, miRNA-3A intermediate MFE, 5A & miRNA-3A cofold MFE, ΔMFE (MFE of miRNA-3A subtracted from of MFE of the 5A & miRNA-3A cofold complex), and predicted ligation loop length - and observed ligation bias for standard Illumina TruSeq (5A), baseline 5A3, prototypical 5A3C6, and 5A3C3 adapters. 88x synthetic miRNA panel was used for this set of experiments.

B) Scatter plots depicting linear regressions and Spearman correlation statistics (ρ and p-values) for ΔMFE (MFE of miRNA-3A subtracted from of MFE of the 5A & miRNA-3A cofold complex) in kcal/mol versus log2-transformed ratio of observed to expected read counts (Bias) across the four evaluated adapter variants.

C) Stacked bar charts showing the relative percentage distribution of empirically determined bimolecular cofold types formed between the respective 5' adapters and the target miRNA library pool. This analysis did not enforce the intermolecular interaction restriction used for ΔMFE calculations.

D) Boxplots depicting ligation bias, expressed as the log2-transformed ratio of observed to expected read counts, grouped across investigated cofold types. Results show no relationship between cofold type and empirically observed bias. Kruskal-Wallis test was used to compute possible statistical significance of relationship between cofold type and empirical bias, and the resulting H-statistic and p-value can be found inside parenthesis located after each legend label.

## S4A

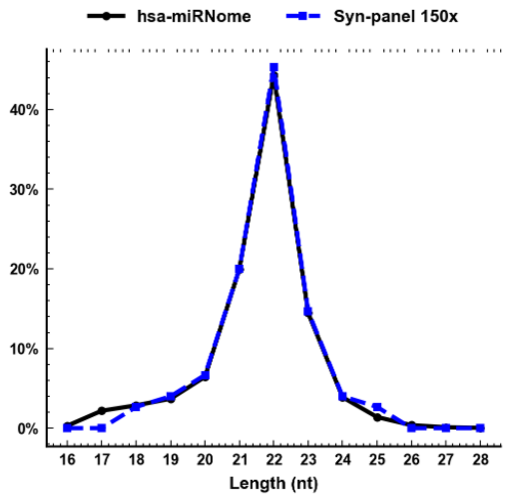


## S4B

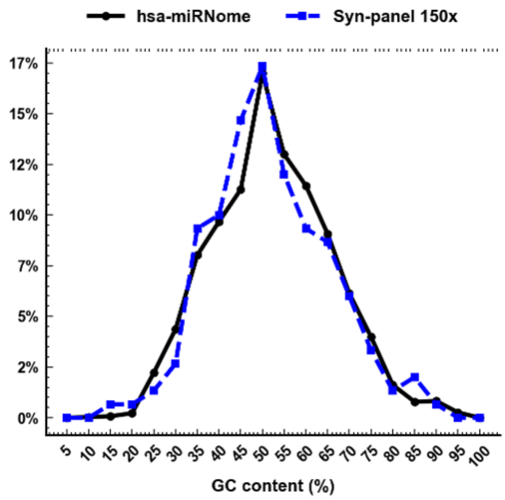


## S4C

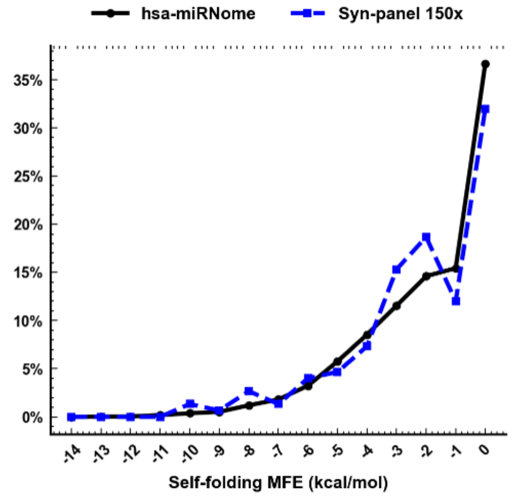


## S4D

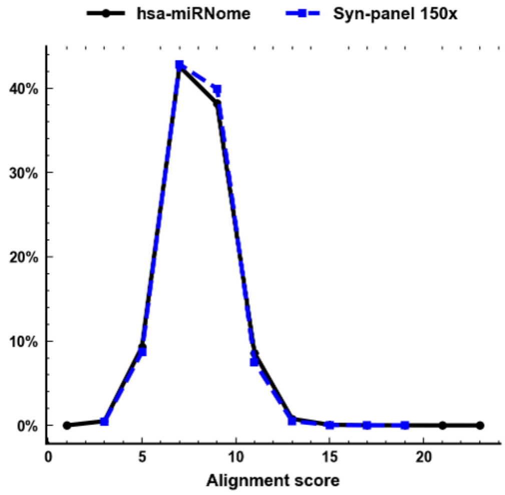

**Supplementary Figure S4. Frequency distribution profiles characterizing the 150x synthetic human miRNA panel.**

A) Line plot comparing the frequency distribution of sequence length (nucleotides, nt) between the total annotated natural human miRNome (miRBase v22.1) and the curated 150x synthetic human miRNA panel (Syn-panel 150x).

B) Line plot comparing the frequency distribution of GC content (%) between the total human miRNome and the curated 150x synthetic human miRNA panel.

C) Line plot comparing the frequency distribution of native secondary structure stability, expressed as miRNA minimum self-folding energy (kcal/mol), between the total human miRNome and the curated 150x synthetic human miRNA panel.

D) Line plot comparing the frequency distribution of predicted inter-sequence alignment scores (obtained by using BioPython's Aligner for all pairwise heterodimer interactions) between the total human miRNome and the curated 150x synthetic human miRNA panel.

S5A

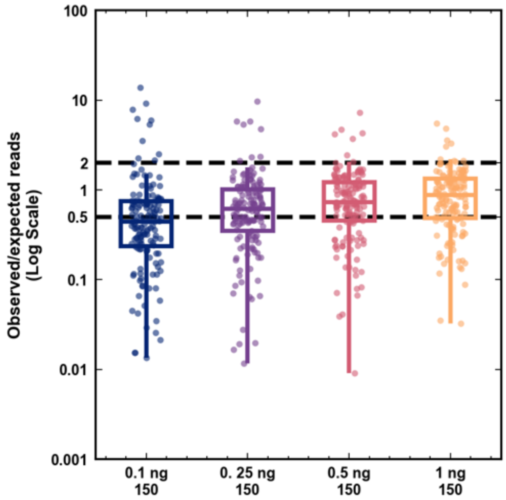


S5B

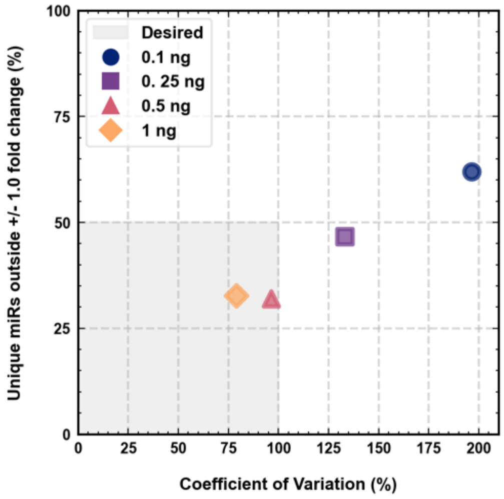


S5C

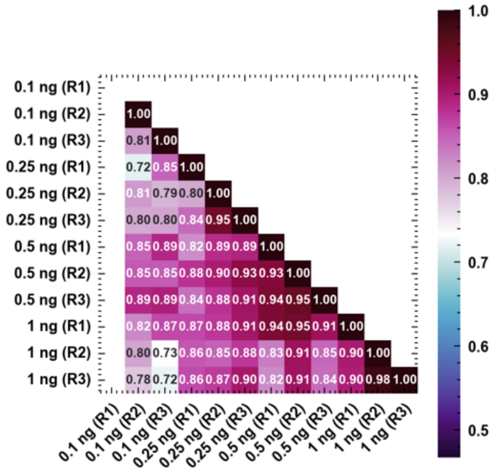


S5D

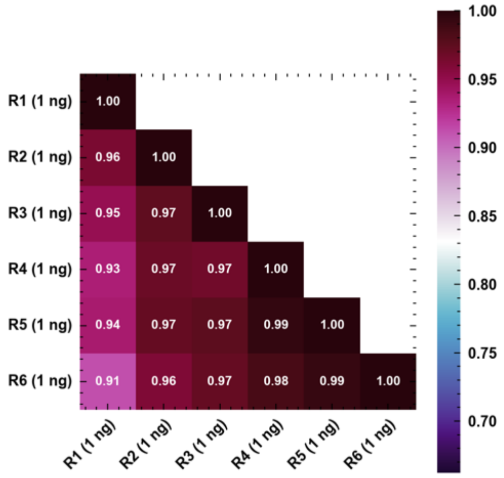

**Supplementary Figure S5. Sensitivity, detection limits, and technical reproducibility of the MDX protocol.**

A) Boxplots depicting ligation bias (average of 3 technical replicates), expressed as the log2-transformed ratio of observed to expected read counts, across ultra-low-input total RNA equivalents ranging from 0.1 ng to 1 ng using the 150x synthetic miRNA reference panel. Numbers underneath group labels indicate number of unique miRNAs detected for individual groups. Horizontal dashed lines define the +/- 1 log2 fold change low-bias range.

B) Scatter plot of above summarizing benchmarking performance by plotting the percentage of unique miRNAs residing outside a +/- 1 log2 fold change window against the overall Coefficient of Variation (CV, %) across input amounts, illustrating the shift in ligation dynamics below 0.5 ng.

C) Heatmap of log-log Pearson correlation coefficients calculated across 3 independent low-input technical replicates (R1, R2, R3) to evaluate assay consistency across 4 input levels.

D) Heatmap of log-log Pearson correlation coefficients evaluated across 6 independent technical replicates (R1–R6) prepared using 1 ng of total RNA equivalent input, confirming high protocol robustness and technical reproducibility.

| Supplementary Table 1 - miRNA composition of 150x and 88x synthetic panels used. | |
|---|---|
| 150x panel | |
| **miRNA** | **Sequence (5' - 3')** |
| MIR4770 | /5' Phos/rUrGrArGrArUrGrArCrArCrUrGrUrArGrCrU |
| MIR320E | /5' Phos/rArArArGrCrUrGrGrGrUrUrGrArGrArArGrG |
| MIR4314 | /5' Phos/rCrUrCrUrGrGrGrArArArUrGrGrGrArCrArG |
| MIR3665 | /5' Phos/rArGrCrArGrGrUrGrCrGrGrGrGrCrGrGrCrG |
| MIR302D-1 | /5' Phos/rArArArArGrCrUrGrGrGrUrUrGrArGrArGrGrA |
| MIR563 | /5' Phos/rArGrGrUrUrGrArCrArUrArCrGrUrUrUrCrCrC |
| MIR4736 | /5' Phos/rArGrGrCrArGrGrUrUrArUrCrUrGrGrGrCrUrG |
| MIR4330 | /5' Phos/rCrCrUrCrArGrArUrCrArGrArGrCrCrUrUrGrC |
| MIR575 | /5' Phos/rGrArGrCrCrArGrUrUrGrGrArCrArGrGrArGrC |
| MIR6127 | /5' Phos/rUrGrArGrGrGrArGrUrGrGrGrUrGrGrGrArGrG |
| MIR511-3p | /5' Phos/rArArUrGrUrGrUrArGrCrArArArArGrArCrArGrA |
| MIR599 | /5' Phos/rGrUrUrGrUrGrUrCrArGrUrUrUrArUrCrArArArC |
| MIR591 | /5' Phos/rArGrArCrCrArUrGrGrGrUrUrCrUrCrArUrUrGrU |
| MIR320C1 | /5' Phos/rArArArArGrCrUrGrGrGrUrUrGrArGrArGrGrGrU |
| MIR147A | /5' Phos/rGrUrGrUrGrUrGrGrArArArUrGrCrUrUrCrUrGrC |
| MIR4429 | /5' Phos/rArArArArGrCrUrGrGrGrCrUrGrArGrArGrGrCrG |
| MIR326 | /5' Phos/rCrCrUrCrUrGrGrGrCrCrCrUrUrCrCrUrCrCrArG |
| MIR937 | /5' Phos/rGrUrGrArGrUrCrArGrGrGrUrGrGrGrGrCrUrGrG |
| MIR6784 | /5' Phos/rGrCrCrGrGrGrGrCrUrUrUrGrGrGrUrGrArGrGrG |
| MIR1199 | /5' Phos/rCrCrUrGrArGrCrCrCrGrGrGrCrCrGrCrGrCrArG |
| MIR590-3p | /5' Phos/rUrArArUrUrUrUrArUrGrUrArUrArArGrCrUrArGrU |
| MIR548BC | /5' Phos/rArArArArArCrUrGrUrGrArUrUrArCrUrUrUrUrGrC |
| MIR4735-3p | /5' Phos/rArArArGrGrUrGrCrUrCrArArArUrUrArGrArCrArU |
| MIR607 | /5' Phos/rGrUrUrCrArArArUrCrCrArGrArUrCrUrArUrArArC |
| MIR26B | /5' Phos/rUrUrCrArArGrUrArArUrUrCrArGrGrArUrArGrGrU |
| MIR218-2 | /5' Phos/rUrUrGrUrGrCrUrUrGrArUrCrUrArArCrCrArUrGrU |
| MIR297 | /5' Phos/rArUrGrUrArUrGrUrGrUrGrCrArUrGrUrGrCrArUrG |
| MIR892C | /5' Phos/rUrArUrUrCrArGrArArArGrGrUrGrCrCrArGrUrCrA |
| MIR411 | /5' Phos/rUrArGrUrArGrArCrCrGrUrArUrArGrCrGrUrArCrG |
| MIR610 | /5' Phos/rUrGrArGrCrUrArArArUrGrUrGrUrGrCrUrGrGrGrA |
| MIR379 | /5' Phos/rUrGrGrUrArGrArCrUrArUrGrGrArArCrGrUrArGrG |
| MIR378H | /5' Phos/rArCrUrGrGrArCrUrUrGrGrUrGrUrCrArGrArUrGrG |
| MIR31 | /5' Phos/rArGrGrCrArArGrArUrGrCrUrGrGrCrArUrArGrCrU |
| MIR323A | /5' Phos/rCrArCrArUrUrArCrArCrGrGrUrCrGrArCrCrUrCrU |
| MIR595 | /5' Phos/rGrArArGrUrGrUrGrCrCrGrUrGrGrUrGrUrGrUrCrU |
| MIR647 | /5' Phos/rGrUrGrGrCrUrGrCrArCrUrCrArCrUrUrCrCrUrUrC |
| MIR140 | /5' Phos/rUrArCrCrArCrArGrGrGrUrArGrArArCrCrArCrGrG |
| MIR4717-3p | /5' Phos/rArCrArCrArUrGrGrGrUrGrGrCrUrGrUrGrGrCrCrU |
| MIR188 | /5' Phos/rCrArUrCrCrCrUrUrGrCrArUrGrGrUrGrGrArGrGrG |
| MIR181D-3p | /5' Phos/rCrCrArCrCrGrGrGrGrGrArUrGrArArUrGrUrCrArC |
| MIR2116-3p | /5' Phos/rCrCrUrCrCrCrArUrGrCrCrArArGrArArCrUrCrCrC |
| MIR486-3p | /5' Phos/rCrGrGrGrGrCrArGrCrUrCrArGrUrArCrArGrGrArU |

| | |
|---|---|
| MIR204-3p | /5' Phos/rGrCrUrGrGrGrArArGrGrCrArArArGrGrGrArCrGrU |
| MIR631 | /5' Phos/rArGrArCrCrUrGrGrCrCrCrArGrArCrCrUrCrArGrC |
| MIR296 | /5' Phos/rArGrGrGrCrCrCrCrCrCrUrCrArArUrCrCrUrGrU |
| MIR4433 | /5' Phos/rCrGrUrCrCrCrArCrCrCrCrCrArCrUrCrCrUrGrU |
| MIR7847 | /5' Phos/rCrGrUrGrGrArGrGrArCrGrArGrGrArGrGrArGrGrC |
| MIR596 | /5' Phos/rArArGrCrCrUrGrCrCrCrGrGrCrUrCrCrUrCrGrGrG |
| MIR149-3p | /5' Phos/rArGrGrGrArGrGrGrArCrGrGrGrGrCrUrGrUrGrC |
| MIR718 | /5' Phos/rCrUrUrCrCrGrCrCrCrCrGrCrCrGrGrGrCrGrUrCrG |
| MIR545 | /5' Phos/rUrCrArGrUrArArArUrGrUrUrArUrUrArGrArUrGrA |
| MIR190A | /5' Phos/rUrGrArUrArUrGrUrUrUrGrArUrArUrArUrUrArGrGrU |
| MIR98 | /5' Phos/rUrGrArGrGrUrArGrUrArArGrUrUrGrUrArUrUrGrUrU |
| MIR542 | /5' Phos/rUrGrUrGrArCrArGrArUrUrGrArUrArArCrUrGrArArA |
| MIR367 | /5' Phos/rArArUrUrGrCrArCrUrUrUrArGrCrArArUrGrGrUrGrA |
| MIR29A | /5' Phos/rArCrUrGrArUrUrUrCrUrUrUrUrGrGrUrGrUrUrCrArG |
| MIR548AC | /5' Phos/rCrArArArArArCrCrGrGrCrArArUrUrArCrUrUrUrUrG |
| MIR548A1 | /5' Phos/rCrArArArArCrUrGrGrCrArArUrUrArCrUrUrUrUrGrC |
| MIR21 | /5' Phos/rUrArGrCrUrUrArUrCrArGrArCrUrGrArUrGrUrUrGrA |
| MIRLET7A2 | /5' Phos/rUrGrArGrGrUrArGrUrArGrGrUrUrGrUrArUrArGrUrU |
| MIR493 | /5' Phos/rUrUrGrUrArCrArUrGrGrUrArGrGrCrUrUrUrCrArUrU |
| MIR216B | /5' Phos/rArArArUrCrUrCrUrGrCrArGrGrCrArArArUrGrUrGrA |
| MIR520F | /5' Phos/rArArGrUrGrCrUrUrCrCrUrUrUrUrArGrArGrGrGrUrU |
| MIR199B | /5' Phos/rArCrArGrUrArGrUrCrUrGrCrArCrArUrUrGrGrUrUrA |
| MIR548B | /5' Phos/rCrArArGrArArCrCrUrCrArGrUrUrGrCrUrUrUrUrGrU |
| MIR627 | /5' Phos/rGrUrGrArGrUrCrUrCrUrArArGrArArArArGrArGrGrA |
| MIR200A | /5' Phos/rUrArArCrArCrUrGrUrCrUrGrGrUrArArCrGrArUrGrU |
| MIRLET7I | /5' Phos/rUrGrArGrGrUrArGrUrArGrUrUrUrGrUrGrCrUrGrUrU |
| MIR30B | /5' Phos/rUrGrUrArArArCrArUrCrCrUrArCrArCrUrCrArGrCrU |
| MIR513C | /5' Phos/rUrUrCrUrCrArArGrGrArGrGrUrGrUrCrGrUrUrUrArU |
| MIR28 | /5' Phos/rArArGrGrArGrCrUrCrArCrArGrUrCrUrArUrUrGrArG |
| MIR643 | /5' Phos/rArCrUrUrGrUrArUrGrCrUrArGrCrUrCrArGrGrUrArG |
| MIR517A | /5' Phos/rArUrCrGrUrGrCrArUrCrCrCrUrUrUrArGrArGrUrGrU |
| MIR216A | /5' Phos/rUrArArUrCrUrCrArGrCrUrGrGrCrArArCrUrGrUrGrA |
| MIR660 | /5' Phos/rUrArCrCrCrArUrUrGrCrArUrArUrCrGrGrArGrUrUrG |
| MIR16-1 | /5' Phos/rUrArGrCrArGrCrArCrGrUrArArArUrArUrUrGrGrCrG |
| MIR654 | /5' Phos/rUrArUrGrUrCrUrGrCrUrGrArCrCrArUrCrArCrCrUrU |
| MIR126 | /5' Phos/rUrCrGrUrArCrCrGrUrGrArGrUrArArUrArArUrGrCrG |
| MIR494 | /5' Phos/rUrGrArArArCrArUrArCrArCrGrGrGrArArArCrCrUrC |
| MIR4510 | /5' Phos/rUrGrArGrGrGrArGrUrArGrGrArUrGrUrArUrGrGrUrU |
| MIR24-1 | /5' Phos/rUrGrCrCrUrArCrUrGrArGrCrUrGrArUrArUrCrArGrU |
| MIR299 | /5' Phos/rUrGrGrUrUrUrArCrCrGrUrCrCrCrArCrArUrArCrArU |
| MIR320B1 | /5' Phos/rArArArArGrCrUrGrGrGrUrUrGrArGrArGrGrGrCrArA |
| MIR99A | /5' Phos/rArArCrCrCrGrUrArGrArUrCrCrGrArUrCrUrUrGrUrG |
| MIR512-1 | /5' Phos/rArArGrUrGrCrUrGrUrCrArUrArGrCrUrGrArGrGrUrC |
| MIR617 | /5' Phos/rArGrArCrUrUrCrCrCrArUrUrUrGrArArGrGrUrGrGrC |
| MIR510 | /5' Phos/rUrArCrUrCrArGrGrArGrArGrUrGrGrCrArArUrCrArC |
| MIR125B | /5' Phos/rUrCrCrCrUrGrArGrArCrCrCrUrArArCrUrUrGrUrGrA |

| | |
|---|---|
| MIR509-3 | /5' Phos/rUrGrArUrUrGrGrUrArCrGrUrCrUrGrUrGrGrGrUrArG |
| MIR184 | /5' Phos/rUrGrGrArCrGrGrArGrArArCrUrGrArUrArArGrGrGrU |
| MIR34A | /5' Phos/rUrGrGrCrArGrUrGrUrCrUrUrArGrCrUrGrGrUrUrGrU |
| MIR380 | /5' Phos/rUrGrGrUrUrGrArCrCrArUrArGrArArCrArUrGrCrGrC |
| MIR133B | /5' Phos/rUrUrUrGrGrUrCrCrCrCrUrUrCrArArCrCrArGrCrUrA |
| MIR504 | /5' Phos/rArGrArCrCrCrUrGrGrUrCrUrGrCrArCrUrCrUrArUrC |
| MIR500A | /5' Phos/rArUrGrCrArCrCrUrGrGrGrCrArArGrGrArUrUrCrUrG |
| MIR218-1-3p | /5' Phos/rArUrGrGrUrUrCrCrGrUrCrArArGrCrArCrCrArUrGrG |
| MIR124 | /5' Phos/rCrGrUrGrUrUrCrArCrArGrCrGrGrArCrCrUrUrGrArU |
| MIR518E | /5' Phos/rCrUrCrUrArGrArGrGrGrArArGrCrGrCrUrUrUrCrUrG |
| MIR30C1-3p | /5' Phos/rCrUrGrGrGrArGrArGrGrGrUrUrGrUrUrUrArCrUrCrC |
| MIR205 | /5' Phos/rUrCrCrUrUrCrArUrUrCrCrArCrCrGrGrArGrUrCrUrG |
| MIR150 | /5' Phos/rUrCrUrCrCrCrArArCrCrCrUrUrGrUrArCrCrArGrUrG |
| MIR891A | /5' Phos/rUrGrCrArArCrGrArArCrCrUrGrArGrCrCrArCrUrGrA |
| MIR541-3p | /5' Phos/rUrGrGrUrGrGrGrCrArCrArGrArArUrCrUrGrGrArCrU |
| MIR375 | /5' Phos/rUrUrUrGrUrUrCrGrUrUrCrGrGrCrUrCrGrCrGrUrGrA |
| MIR193B | /5' Phos/rCrGrGrGrGrUrUrUrUrGrArGrGrGrCrGrArGrArUrGrA |
| MIR23A | /5' Phos/rGrGrGrGrUrUrCrCrUrGrGrGrGrArUrGrGrGrArUrUrU |
| MIR18B-3p | /5' Phos/rUrGrCrCrCrUrArArArUrGrCrCrCrCrUrUrCrUrGrGrC |
| MIR134 | /5' Phos/rUrGrUrGrArCrUrGrGrUrUrGrArCrCrArGrArGrGrGrG |
| MIR99B | /5' Phos/rCrArCrCrCrGrUrArGrArArCrCrGrArCrCrUrUrGrCrG |
| MIR711 | /5' Phos/rGrGrGrArCrCrCrArGrGrGrArGrArGrArCrGrUrArArG |
| MIR486 | /5' Phos/rUrCrCrUrGrUrArCrUrGrArGrCrUrGrCrCrCrCrGrArG |
| MIR654 | /5' Phos/rUrGrGrUrGrGrGrCrCrGrCrArGrArArCrArUrGrUrGrC |
| MIR2110 | /5' Phos/rUrUrGrGrGrGrArArArCrGrGrCrCrGrCrUrGrArGrUrG |
| MIR766 | /5' Phos/rArCrUrCrCrArGrCrCrCrCrArCrArGrCrCrUrCrArGrC |
| MIR370 | /5' Phos/rGrCrCrUrGrCrUrGrGrGrGrUrGrGrArArCrCrUrGrGrU |
| MIR187 | /5' Phos/rGrGrCrUrArCrArArCrArCrArGrGrArCrCrCrGrGrGrC |
| MIR92B | /5' Phos/rArGrGrGrArCrGrGrGrArCrGrCrGrGrUrGrCrArGrUrG |
| MIR663 | /5' Phos/rArGrGrCrGrGrGrGrCrGrCrCrGrCrGrGrGrArCrCrGrC |
| MIR4668 | /5' Phos/rArGrGrGrArArArArArArArArArArGrGrArUrUrUrGrUrC |
| MIR29B-3p | /5' Phos/rUrArGrCrArCrCrArUrUrUrGrArArArUrCrArGrUrGrUrU |
| MIR135B | /5' Phos/rUrArUrGrGrCrUrUrUrUrCrArUrUrCrCrUrArUrGrUrGrA |
| MIR548U | /5' Phos/rCrArArArGrArCrUrGrCrArArUrUrArCrUrUrUrUrGrCrG |
| MIR301B | /5' Phos/rCrArGrUrGrCrArArUrGrArUrArUrUrGrUrCrArArArGrC |
| MIR181D | /5' Phos/rArArCrArUrUrCrArUrUrGrUrUrGrUrCrGrGrUrGrGrGrU |
| MIR10B | /5' Phos/rUrArCrCrCrUrGrUrArGrArArCrCrGrArArUrUrUrGrUrG |
| MIR217 | /5' Phos/rUrArCrUrGrCrArUrCrArGrGrArArCrUrGrArUrUrGrGrA |
| MIR146B | /5' Phos/rUrGrArGrArArCrUrGrArArUrUrCrCrArUrArGrGrCrUrG |
| MIR181A1 | /5' Phos/rArArCrArUrUrCrArArCrGrCrUrGrUrCrGrGrUrGrArGrU |
| MIR425 | /5' Phos/rArArUrGrArCrArCrGrArUrCrArCrUrCrCrCrGrUrUrGrA |
| MIR574 | /5' Phos/rUrGrArGrUrGrUrGrUrGrUrGrUrGrUrGrArGrUrGrUrGrU |
| MIR191 | /5' Phos/rCrArArCrGrGrArArUrCrCrCrArArArArGrCrArGrCrUrG |
| MIR767 | /5' Phos/rUrGrCrArCrCrArUrGrGrUrUrGrUrCrUrGrArGrCrArUrG |
| MIR371B-3p | /5' Phos/rArArGrUrGrCrCrCrCrCrArCrArGrUrUrUrGrArGrUrGrC |
| MIR512-2 | /5' Phos/rCrArCrUrCrArGrCrCrUrUrGrArGrGrGrCrArCrUrUrUrC |

| | |
|---|---|
| MIR6837-3p | /5' Phos/rCrCrUrUrCrArCrUrGrUrGrArCrUrCrUrGrCrUrGrCrArG |
| MIR3188 | /5' Phos/rArGrArGrGrCrUrUrUrGrUrGrCrGrGrArUrArCrGrGrGrG |
| MIR128-1 | /5' Phos/rCrGrGrGrGrCrCrGrUrArGrCrArCrUrGrUrCrUrGrArGrA |
| MIR339 | /5' Phos/rUrCrCrCrUrGrUrCrCrUrCrCrArGrGrArGrCrUrCrArCrG |
| MIR611 | /5' Phos/rGrCrGrArGrGrArCrCrCrCrUrCrGrGrGrGrUrCrUrGrArC |
| MIR602 | /5' Phos/rGrArCrArCrGrGrGrCrGrArCrArGrCrUrGrCrGrGrCrCrC |
| MIR1277 | /5' Phos/rArArArUrArUrArUrArUrArUrArUrArUrGrUrArCrGrUrArU |
| MIR3613-3p | /5' Phos/rArCrArArArArArArArArArArGrCrCrCrArArCrCrCrUrUrC |
| MIR8065 | /5' Phos/rUrGrUrArGrGrArArCrArGrUrUrGrArArUrUrUrUrGrGrCrU |
| MIR182 | /5' Phos/rUrUrUrGrGrCrArArUrGrGrUrArGrArArCrUrCrArCrArCrU |
| MIR637 | /5' Phos/rArCrUrGrGrGrGrGrCrUrUrUrCrGrGrGrCrUrCrUrGrCrGrU |
| MIR939 | /5' Phos/rUrGrGrGrGrArGrCrUrGrArGrGrCrUrCrUrGrGrGrGrGrUrG |
| MIR378C | /5' Phos/rArCrUrGrGrArCrUrUrGrGrArGrUrCrArGrArArGrArGrUrGrG |
| MIR608 | /5' Phos/rArGrGrGrGrUrGrGrUrGrUrUrGrGrGrArCrArGrCrUrCrCrGrU |
| MIR658 | /5' Phos/rGrGrCrGrGrArGrGrGrArArGrUrArGrGrUrCrCrGrUrUrGrGrU |
| MIR638 | /5' Phos/rArGrGrGrArUrCrGrCrGrGrGrCrGrGrGrUrGrGrCrGrGrCrCrU |
| **88x panel** | |
| **miRNA** | **Sequence (5' - 3')** |
| MIR3168 | /5' Phos/rGrArGrUrUrCrUrArCrArGrUrCrArGrArC |
| MIR4770 | /5' Phos/rUrGrArGrArUrGrArCrArCrUrGrUrArGrCrU |
| MIR320E | /5' Phos/rArArArGrCrUrGrGrGrUrUrGrArGrArArGrG |
| MIR575 | /5' Phos/rGrArGrCrCrArGrUrUrGrGrArCrArGrGrArGrC |
| MIR599 | /5' Phos/rGrUrUrGrUrGrUrCrArGrUrUrUrArUrCrArArArC |
| MIR591 | /5' Phos/rArGrArCrCrArUrGrGrGrUrUrCrUrCrArUrUrGrU |
| MIR320C1 | /5' Phos/rArArArArGrCrUrGrGrGrUrUrGrArGrArGrGrGrU |
| MIR147A | /5' Phos/rGrUrGrUrGrUrGrGrArArArUrGrCrUrUrCrUrGrC |
| MIR4429 | /5' Phos/rArArArArGrCrUrGrGrGrCrUrGrArGrArGrGrCrG |
| MIR326 | /5' Phos/rCrCrUrCrUrGrGrGrCrCrCrUrUrCrCrUrCrCrArG |
| MIR607 | /5' Phos/rGrUrUrCrArArArUrCrCrArGrArUrCrUrArUrArArC |
| MIR26B | /5' Phos/rUrUrCrArArGrUrArArUrUrCrArGrGrArUrArGrGrU |
| MIR218-2 | /5' Phos/rUrUrGrUrGrCrUrUrGrArUrCrUrArArCrCrArUrGrU |
| MIR297 | /5' Phos/rArUrGrUrArUrGrUrGrUrGrCrArUrGrUrGrCrArUrG |
| MIR892C | /5' Phos/rUrArUrUrCrArGrArArArGrGrUrGrCrCrArGrUrCrA |
| MIR411 | /5' Phos/rUrArGrUrArGrArCrCrGrUrArUrArGrCrGrUrArCrG |
| MIR610 | /5' Phos/rUrGrArGrCrUrArArArUrGrUrGrUrGrCrUrGrGrGrA |
| MIR379 | /5' Phos/rUrGrGrUrArGrArCrUrArUrGrGrArArCrGrUrArGrG |
| MIR378H | /5' Phos/rArCrUrGrGrArCrUrUrGrGrUrGrUrCrArGrArUrGrG |
| MIR323A | /5' Phos/rCrArCrArUrUrArCrArCrGrGrUrCrGrArCrCrUrCrU |
| MIR595 | /5' Phos/rGrArArGrUrGrUrGrCrCrGrUrGrGrUrGrUrGrUrCrU |
| MIR647 | /5' Phos/rGrUrGrGrCrUrGrCrArCrUrCrArCrUrUrCrCrUrUrC |
| MIR140 | /5' Phos/rUrArCrCrArCrArGrGrGrUrArGrArArCrCrArCrGrG |
| MIR7847 | /5' Phos/rCrGrUrGrGrArGrGrArCrGrArGrGrArGrGrArGrGrC |
| MIR718 | /5' Phos/rCrUrUrCrCrGrCrCrCrCrGrCrCrGrGrGrCrGrUrCrG |
| MIR98 | /5' Phos/rUrGrArGrGrUrArGrUrArArGrUrUrGrUrArUrUrGrUrU |
| MIR542 | /5' Phos/rUrGrUrGrArCrArGrArUrUrGrArUrArArCrUrGrArArA |
| MIR367 | /5' Phos/rArArUrUrGrCrArCrUrUrUrArGrCrArArUrGrGrUrGrA |

| | |
|---|---|
| MIR548AC | /5' Phos/rCrArArArArArCrCrGrGrCrArArUrUrArCrUrUrUrG |
| MIR548A1 | /5' Phos/rCrArArArArCrUrGrGrCrArArUrUrArCrUrUrUrUrGrC |
| MIR21 | /5' Phos/rUrArGrCrUrUrArUrCrArGrArCrUrGrArUrGrUrUrGrA |
| MIRLET7A2 | /5' Phos/rUrGrArGrGrUrArGrUrArGrGrUrUrGrUrArUrArGrUrU |
| MIR493 | /5' Phos/rUrUrGrUrArCrArUrGrGrUrArGrGrCrUrUrUrCrArUrU |
| MIR216B | /5' Phos/rArArArUrCrUrCrUrGrCrArGrGrCrArArArUrGrUrGrA |
| MIR520F | /5' Phos/rArArGrUrGrCrUrUrCrCrUrUrUrUrArGrArGrGrGrUrU |
| MIR199B | /5' Phos/rArCrArGrUrArGrUrCrUrGrCrArCrArUrUrGrGrUrUrA |
| MIR548B | /5' Phos/rCrArArGrArArCrCrUrCrArGrUrUrGrCrUrUrUrUrGrU |
| MIR627 | /5' Phos/rGrUrGrArGrUrCrUrCrUrArArGrArArArArGrArGrGrA |
| MIR200A | /5' Phos/rUrArArCrArCrUrGrUrCrUrGrGrUrArArCrGrArUrGrU |
| MIRLET7I | /5' Phos/rUrGrArGrGrUrArGrUrArGrUrUrUrGrUrGrCrUrGrUrU |
| MIR30B | /5' Phos/rUrGrUrArArArCrArUrCrCrUrArCrArCrUrCrArGrCrU |
| MIR513C | /5' Phos/rUrUrCrUrCrArArGrGrArGrGrUrGrUrCrGrUrUrUrArU |
| MIR28 | /5' Phos/rArArGrGrArGrCrUrCrArCrArGrUrCrUrArUrUrGrArG |
| MIR643 | /5' Phos/rArCrUrUrGrUrArUrGrCrUrArGrCrUrCrArGrGrUrArG |
| MIR517A | /5' Phos/rArUrCrGrUrGrCrArUrCrCrCrUrUrUrArGrArGrUrGrU |
| MIR216A | /5' Phos/rUrArArUrCrUrCrArGrCrUrGrGrCrArArCrUrGrUrGrA |
| MIR660 | /5' Phos/rUrArCrCrCrArUrUrGrCrArUrArUrCrGrGrArGrUrUrG |
| MIR654 | /5' Phos/rUrArUrGrUrCrUrGrCrUrGrArCrCrArUrCrArCrCrUrU |
| MIR126 | /5' Phos/rUrCrGrUrArCrCrGrUrGrArGrUrArArUrArArUrGrCrG |
| MIR494 | /5' Phos/rUrGrArArArCrArUrArCrArCrGrGrGrArArArCrCrUrC |
| MIR4510 | /5' Phos/rUrGrArGrGrGrArGrUrArGrGrArUrGrUrArUrGrGrUrU |
| MIR299 | /5' Phos/rUrGrGrUrUrUrArCrCrGrUrCrCrCrArCrArUrArCrArU |
| MIR320B1 | /5' Phos/rArArArArGrCrUrGrGrGrUrUrGrArGrArGrGrGrCrArA |
| MIR99A | /5' Phos/rArArCrCrCrGrUrArGrArUrCrCrGrArUrCrUrUrGrUrG |
| MIR512-1 | /5' Phos/rArArGrUrGrCrUrGrUrCrArUrArGrCrUrGrArGrGrUrC |
| MIR617 | /5' Phos/rArGrArCrUrUrCrCrCrArUrUrUrGrArArGrGrUrGrGrC |
| MIR510 | /5' Phos/rUrArCrUrCrArGrGrArGrArGrUrGrGrCrArArUrCrArC |
| MIR509-3 | /5' Phos/rUrGrArUrUrGrGrUrArCrGrUrCrUrGrUrGrGrGrUrArG |
| MIR184 | /5' Phos/rUrGrGrArCrGrGrArGrArArCrUrGrArUrArArGrGrGrU |
| MIR380 | /5' Phos/rUrGrGrUrUrGrArCrCrArUrArGrArArCrArUrGrCrGrC |
| MIR133B | /5' Phos/rUrUrUrGrGrUrCrCrCrCrUrUrCrArArCrCrArGrCrUrA |
| MIR504 | /5' Phos/rArGrArCrCrCrUrGrGrUrCrUrGrCrArCrUrCrUrArUrC |
| MIR500A | /5' Phos/rArUrGrCrArCrCrUrGrGrGrCrArArGrGrArUrUrCrUrG |
| MIR518E | /5' Phos/rCrUrCrUrArGrArGrGrGrArArGrCrGrCrUrUrUrCrUrG |
| MIR205 | /5' Phos/rUrCrCrUrUrCrArUrUrCrCrArCrCrGrGrArGrUrCrUrG |
| MIR150 | /5' Phos/rUrCrUrCrCrCrArArCrCrCrUrUrGrUrArCrCrArGrUrG |
| MIR891A | /5' Phos/rUrGrCrArArCrGrArArCrCrUrGrArGrCrCrArCrUrGrA |
| MIR375 | /5' Phos/rUrUrUrGrUrUrCrGrUrUrCrGrGrCrUrCrGrCrGrUrGrA |
| MIR134 | /5' Phos/rUrGrUrGrArCrUrGrGrUrUrGrArCrCrArGrArGrGrGrG |
| MIR99B | /5' Phos/rCrArCrCrCrGrUrArGrArArCrCrGrArCrCrUrUrGrCrG |
| MIR766 | /5' Phos/rArCrUrCrCrArGrCrCrCrCrArCrArGrCrCrUrCrArGrC |
| MIR370 | /5' Phos/rGrCrCrUrGrCrUrGrGrGrGrUrGrGrArArCrCrUrGrGrU |
| MIR548U | /5' Phos/rCrArArArGrArCrUrGrCrArArUrUrArCrUrUrUrUrGrCrG |
| MIR301B | /5' Phos/rCrArGrUrGrCrArArUrGrArUrArUrUrGrUrCrArArArGrC |

| | |
|---|---|
| MIR181D | /5' Phos/rArArCrArUrUrCrArUrUrGrUrUrGrUrCrGrGrUrGrGrGrU |
| MIR10B | /5' Phos/rUrArCrCrCrUrGrUrArGrArArCrCrGrArArUrUrUrGrUrG |
| MIR217 | /5' Phos/rUrArCrUrGrCrArUrCrArGrGrArArCrUrGrArUrUrGrGrA |
| MIR146B | /5' Phos/rUrGrArGrArArCrUrGrArArUrUrCrCrArUrArGrGrCrUrG |
| MIR181A1 | /5' Phos/rArArCrArUrUrCrArArCrGrCrUrGrUrCrGrGrUrGrArGrU |
| MIR425 | /5' Phos/rArArUrGrArCrArCrGrArUrCrArCrUrCrCrCrGrUrUrGrA |
| MIR191 | /5' Phos/rCrArArCrGrGrArArUrCrCrCrArArArArGrCrArGrCrUrG |
| MIR767 | /5' Phos/rUrGrCrArCrCrArUrGrGrUrUrGrUrCrUrGrArGrCrArUrG |
| MIR339 | /5' Phos/rUrCrCrCrUrGrUrCrCrUrCrCrArGrGrArGrCrUrCrArCrG |
| MIR602 | /5' Phos/rGrArCrArCrGrGrGrCrGrArCrArGrCrUrGrCrGrGrCrCrC |
| MIR8065 | /5' Phos/rUrGrUrArGrGrArArCrArGrUrUrGrArArUrUrUrUrGrGrCrU |
| MIR182 | /5' Phos/rUrUrUrGrGrCrArArUrGrGrUrArGrArArCrUrCrArCrArCrU |
| MIR378C | /5' Phos/rArCrUrGrGrArCrUrUrGrGrArGrUrCrArGrArArGrArGrUrGrG |
| MIR608 | /5' Phos/rArGrGrGrGrUrGrGrUrGrUrUrGrGrGrArCrArGrCrUrCrCrGrU |

| Supplementary Table 2 - List of all adapters, FLRs, PCR primers, and auxiliary oligos used. | | |
|---|---|---|
| **Name** | **Note** | **Sequence (5' - 3')** |
| 5A3 | 5' adapter | rGrUrUrCrArGrArGrUrUrCrUrArCrArGrUrCrCrGrArCrGrArUrCrNrN |
| 5A3C1 | 5' adapter | rGrUrUrCrArGrArGrUrUrCrUrArCrArGrUrCrCrGrArCrGrArUrCrArArCrCrCrGrArGrNrN |
| 5A3C2 | 5' adapter | rGrUrUrCrArGrArGrUrUrCrUrArCrArGrUrCrCrGrArCrGrArUrCrArUrUrGrGrCrArCrArC rCrCrGrArGrNrN |
| 5A3C3 | 5' adapter | rUrUrGrGrCrArCrGrUrUrCrArGrArGrUrUrCrUrArCrArGrUrCrCrGrArCrGrArUrCrArArC rCrCrGrArGrNrN |
| 5A3C4 | 5' adapter | rGrUrUrCrArGrArGrUrUrCrUrArCrArGrUrCrCrGrArCrGrArUrCrArUrUrGrGrCrArCrNrN |
| 5A3C5 | 5' adapter | rUrUrGrGrCrArCrArArCrCrCrGrArGrGrUrUrCrArGrArGrUrUrCrUrArCrArGrUrCrCrGrA rCrGrArUrCrNrN |
| 5A3C6 | 5' adapter | rArCrCrCrGrArGrGrUrUrCrArGrArGrUrUrCrUrArCrArGrUrCrCrGrArCrGrArUrCrNrN |
| 5A3C7 | 5' adapter | rUrUrGrGrCrArCrGrUrUrCrArGrArGrUrUrCrUrArCrArGrUrCrCrGrArCrGrArUrCrNrN |
| 5A3C6U | 5' adapter | rArCrCrCrGrArGrGrUrUrCrArGrArGrUrUrCrUrArCrArGrUrCrCrGrArCrGrArUrCrNrNrN rArNrNrNrNrArNrNrNrGrArUrCrNrN |
| 3AB1 | 3' adapter | /5'-rApp/NNTGACGTTGGAATTCTCGGG/T-C6-AF488/GCCAAGG/C6-AF488/ |
| dC11 | carrier oligo | TCGAAGTATTC |
| FLR-25 | fluorescent ruler | /5'-5' dC/CTGAGTAACTTAGGTATCCCTAAGTCGCGTGAGGCGTCTCACGCAA/T-C6-AF488/GAAACT/C6-AF488/ |
| FLR-18 | fluorescent ruler | /5'-5' dC/ACTTAGGTATCCCTAAGTCGCGTGAGGCGTCTCACGCAA/T-C6-AF488/GAAACT/C6-AF488/ |
| L30 | ladder oligo | CTAAGGAAACTACAAGAC/T-C6-AF488/AACGCTCCAGT/C6-AF488/ |
| L40 | ladder oligo | CCAACGGCAGCTAGGAATC/T-C6-AF488/ATGAGGATAGTCAAAGGATT/C6-AF488/ |
| L50 | ladder oligo | TGCTCGGTAATTTATGCTCTGGT/T-C6-AF488/ATCTAACTATTCCCATTGTCTAAAGA/C6-AF488/ |
| L60 | ladder oligo | GGACCTATGTTTTCTGCTTGTCGATCCA/T-C6-AF488/AGAAGTCATATCAACGCGCAGGATTTGTGTC/C6-AF488/ |
| L70 | ladder oligo | CTAAGGAAACTACAAGACTAACGCTCCAG/T-C6-AF488/CCAACGGCAGCTAGGAATCTATGAGGATAGTCAAAGGATT/C6-AF488/ |
| L80 | ladder oligo | CTAAGGAAACTACAAGACTAACGCTCCAG/T-C6-AF488/TGCTCGGTAATTTATGCTCTGGTTATCTAACTATTCCCATTGTCTAAAGA/C6-AF488/ |
| RTP | RT primer | GCCTTGGCACCCGAGAATTCCA |
| 5AP1 | 5' PCR primer | AATGATACGGCGACCACCGAGATCTACACGTTCAGAGTTCTACAGTCCGA |
| RPI-ex10-1 | 3' PCR primer (indexed) | CAAGCAGAAGACGGCATACGAGATACCGGCTCAGGTGACTGGAGTTCCTTGGCACCCGAGAATTCCA |
| RPI-ex10-2 | 3' PCR primer (indexed) | CAAGCAGAAGACGGCATACGAGATCAAGGCTATCGTGACTGGAGTTCCTTGGCACCCGAGAATTCCA |
| RPI-ex10-3 | 3' PCR primer (indexed) | CAAGCAGAAGACGGCATACGAGATGTGGTTGAAGGTGACTGGAGTTCCTTGGCACCCGAGAATTCCA |
| RPI-ex10-4 | 3' PCR primer (indexed) | CAAGCAGAAGACGGCATACGAGATTCCATAATCCGTGACTGGAGTTCCTTGGCACCCGAGAATTCCA |
| RPI-ex10-5 | 3' PCR primer (indexed) | CAAGCAGAAGACGGCATACGAGATACTGAATAGAGTGACTGGAGTTCCTTGGCACCCGAGAATTCCA |
| RPI-ex10-6 | 3' PCR primer (indexed) | CAAGCAGAAGACGGCATACGAGATCACGTTAGGCGTGACTGGAGTTCCTTGGCACCCGAGAATTCCA |
| RPI-ex10-7 | 3' PCR primer (indexed) | CAAGCAGAAGACGGCATACGAGATGGACGTCTTGGTGACTGGAGTTCCTTGGCACCCGAGAATTCCA |
| RPI-ex10-8 | 3' PCR primer (indexed) | CAAGCAGAAGACGGCATACGAGATTCGTGGTTGAGTGACTGGAGTTCCTTGGCACCCGAGAATTCCA |
| RPI-ex10-10 | 3' PCR primer (indexed) | CAAGCAGAAGACGGCATACGAGATCACTCGCACTGTGACTGGAGTTCCTTGGCACCCGAGAATTCCA |
| RPI-ex10-11 | 3' PCR primer (indexed) | CAAGCAGAAGACGGCATACGAGATGCAGGCTGGAGTGACTGGAGTTCCTTGGCACCCGAGAATTCCA |
| RPI-ex10-12 | 3' PCR primer (indexed) | CAAGCAGAAGACGGCATACGAGATTGAACGCAACGTGACTGGAGTTCCTTGGCACCCGAGAATTCCA |
| RPI-ex10-13 | 3' PCR primer (indexed) | CAAGCAGAAGACGGCATACGAGATATGAGAACCAGTGACTGGAGTTCCTTGGCACCCGAGAATTCCA |
| RPI-ex10-14 | 3' PCR primer (indexed) | CAAGCAGAAGACGGCATACGAGATCGCCATACCTGTGACTGGAGTTCCTTGGCACCCGAGAATTCCA |
| RPI-ex10-15 | 3' PCR primer (indexed) | CAAGCAGAAGACGGCATACGAGATGTTATATGGCGTGACTGGAGTTCCTTGGCACCCGAGAATTCCA |
| RPI-ex10-16 | 3' PCR primer (indexed) | CAAGCAGAAGACGGCATACGAGATTTGGCCAGGTGTGACTGGAGTTCCTTGGCACCCGAGAATTCCA |
| RPI-ex10-18 | 3' PCR primer (indexed) | CAAGCAGAAGACGGCATACGAGATAGCTTACACAGTGACTGGAGTTCCTTGGCACCCGAGAATTCCA |
| RPI-ex10-19 | 3' PCR primer (indexed) | CAAGCAGAAGACGGCATACGAGATATTGACACATGTGACTGGAGTTCCTTGGCACCCGAGAATTCCA |
| RPI-ex10-20 | 3' PCR primer (indexed) | CAAGCAGAAGACGGCATACGAGATGCAACCAGTCGTGACTGGAGTTCCTTGGCACCCGAGAATTCCA |
| RPI-ex10-22 | 3' PCR primer (indexed) | CAAGCAGAAGACGGCATACGAGATCCGGACCACAGTGACTGGAGTTCCTTGGCACCCGAGAATTCCA |

| | | |
|---|---|---|
| RPI-ex10-23 | 3' PCR primer (indexed) | CAAGCAGAAGACGGCATACGAGATATTCATTGCAGTGACTGGAGTTCCTTGGCACCCGAGAATTCCA |
| RPI-ex10-24 | 3' PCR primer (indexed) | CAAGCAGAAGACGGCATACGAGATTCCTTCATAGGTGACTGGAGTTCCTTGGCACCCGAGAATTCCA |
| RPI-ex10-25 | 3' PCR primer (indexed) | CAAGCAGAAGACGGCATACGAGATTGTGATGTATGTGACTGGAGTTCCTTGGCACCCGAGAATTCCA |
| RPI-ex10-26 | 3' PCR primer (indexed) | CAAGCAGAAGACGGCATACGAGATAACATACCTAGTGACTGGAGTTCCTTGGCACCCGAGAATTCCA |
| RPI-ex10-27 | 3' PCR primer (indexed) | CAAGCAGAAGACGGCATACGAGATGGACCTCAATGTGACTGGAGTTCCTTGGCACCCGAGAATTCCA |
| RPI-ex10-28 | 3' PCR primer (indexed) | CAAGCAGAAGACGGCATACGAGATCTCGACTCCTGTGACTGGAGTTCCTTGGCACCCGAGAATTCCA |
| RPI-ex10-29 | 3' PCR primer (indexed) | CAAGCAGAAGACGGCATACGAGATCGTGTATCTTGTGACTGGAGTTCCTTGGCACCCGAGAATTCCA |
| RPI-ex10-30 | 3' PCR primer (indexed) | CAAGCAGAAGACGGCATACGAGATAGTGAGTGAAGTGACTGGAGTTCCTTGGCACCCGAGAATTCCA |
| RPI-ex10-31 | 3' PCR primer (indexed) | CAAGCAGAAGACGGCATACGAGATGAATGCACGAGTGACTGGAGTTCCTTGGCACCCGAGAATTCCA |
| RPI-ex10-32 | 3' PCR primer (indexed) | CAAGCAGAAGACGGCATACGAGATTGGCAATATTGTGACTGGAGTTCCTTGGCACCCGAGAATTCCA |
| RPI-ex10-33 | 3' PCR primer (indexed) | CAAGCAGAAGACGGCATACGAGATCATACTTGAAGTGACTGGAGTTCCTTGGCACCCGAGAATTCCA |
| RPI-ex10-34 | 3' PCR primer (indexed) | CAAGCAGAAGACGGCATACGAGATAACGGAGCGGGTGACTGGAGTTCCTTGGCACCCGAGAATTCCA |
| RPI-ex10-35 | 3' PCR primer (indexed) | CAAGCAGAAGACGGCATACGAGATGACTTAGAAGGTGACTGGAGTTCCTTGGCACCCGAGAATTCCA |
| RPI-ex10-36 | 3' PCR primer (indexed) | CAAGCAGAAGACGGCATACGAGATTCTAGTCTTCGTGACTGGAGTTCCTTGGCACCCGAGAATTCCA |
| RPI-ex10-37 | 3' PCR primer (indexed) | CAAGCAGAAGACGGCATACGAGATCTTGTCTTAAGTGACTGGAGTTCCTTGGCACCCGAGAATTCCA |
| RPI-ex10-38 | 3' PCR primer (indexed) | CAAGCAGAAGACGGCATACGAGATACTTCCTAGCGTGACTGGAGTTCCTTGGCACCCGAGAATTCCA |
| RPI-ex10-39 | 3' PCR primer (indexed) | CAAGCAGAAGACGGCATACGAGATGAAGCGGACCGTGACTGGAGTTCCTTGGCACCCGAGAATTCCA |
| RPI-ex10-40 | 3' PCR primer (indexed) | CAAGCAGAAGACGGCATACGAGATTGCACGAGAAGTGACTGGAGTTCCTTGGCACCCGAGAATTCCA |
| RPI-ex10-41 | 3' PCR primer (indexed) | CAAGCAGAAGACGGCATACGAGATCCTTAGTGCCGTGACTGGAGTTCCTTGGCACCCGAGAATTCCA |
| RPI-ex10-42 | 3' PCR primer (indexed) | CAAGCAGAAGACGGCATACGAGATAAGAGAGGTGGTGACTGGAGTTCCTTGGCACCCGAGAATTCCA |
| RPI-ex10-43 | 3' PCR primer (indexed) | CAAGCAGAAGACGGCATACGAGATGCTCTCGTTGGTGACTGGAGTTCCTTGGCACCCGAGAATTCCA |
| RPI-ex10-44 | 3' PCR primer (indexed) | CAAGCAGAAGACGGCATACGAGATTGTACCGAATGTGACTGGAGTTCCTTGGCACCCGAGAATTCCA |
| RPI-ex10-45 | 3' PCR primer (indexed) | CAAGCAGAAGACGGCATACGAGATGTGCTAGGTGGTGACTGGAGTTCCTTGGCACCCGAGAATTCCA |
| RPI-ex10-46 | 3' PCR primer (indexed) | CAAGCAGAAGACGGCATACGAGATAGCGTGAATGGTGACTGGAGTTCCTTGGCACCCGAGAATTCCA |
| RPI-ex10-47 | 3' PCR primer (indexed) | CAAGCAGAAGACGGCATACGAGATGTGCTATTAAGTGACTGGAGTTCCTTGGCACCCGAGAATTCCA |
| RPI-ex10-48 | 3' PCR primer (indexed) | CAAGCAGAAGACGGCATACGAGATGAGTCTCTCCGTGACTGGAGTTCCTTGGCACCCGAGAATTCCA |
